# Succinate dehydrogenase loss suppresses pyrimidine biosynthesis via succinate-mediated inhibition of aspartate transcarbamylase

**DOI:** 10.1101/2025.02.18.638948

**Authors:** Madeleine L. Hart, David Sokolov, Serwah Danquah, Eric Zheng, Alex D. Doan, Kristian Davidsen, David MacPherson, Lucas B. Sullivan

**Affiliations:** Human Biology Division, Fred Hutchinson Cancer Center, Seattle, WA, 98109, USA

**Keywords:** aspartate, biosensor, metabolism, succinate dehydrogenase, SDH, aspartate transcarbamylase, replication stress, pyrimidines, nucleotides, cancer

## Abstract

Decreased availability of the amino acid aspartate can constrain cell function in diverse biological contexts, but the temporal interplay between aspartate, downstream metabolic changes, and functional effects remains poorly understood. Using an aspartate biosensor and live-cell imaging, we examine the interaction between aspartate abundance and cell proliferation in several models of aspartate limitation. While aspartate deficiencies intuitively interface with proliferation in some contexts, aspartate limitation from succinate dehydrogenase (SDH) inhibition causes strikingly nonintuitive dynamics resulting from an outsized impairment of pyrimidine synthesis. Mechanistically, we find that SDH loss impairs pyrimidine biosynthesis by decreasing aspartate and accumulating succinate, which competitively inhibits mammalian aspartate transcarbamylase (ATCase). This metabolic interaction persists in multiple models of SDH deficiency, causing pyrimidine insufficiency, replication stress, and sensitivity to ATR kinase inhibition. These findings define a novel role for succinate in modulating cellular nucleotide homeostasis, suggest a potential therapeutic vulnerability of SDH-deficient tumors, and demonstrate how cascading metabolic interactions can unfold to impact cell function.

## Introduction

Cellular metabolism is amongst the most dynamic biological processes, with many biochemical reactions operating at sub-second timescales and metabolite pools turning over on the scale of seconds to hours^1^. Metabolic disruptions can therefore trigger rapid and evolving changes that reverberate across the metabolic network and occur with distinct temporal behaviors. The abundance of the amino acid aspartate is highly responsive to metabolic state, since aspartate is predominantly synthesized by cells via mitochondrial metabolism, after which it serves as a precursor for several major anabolic products. Aspartate limitation—the condition where aspartate levels are insufficient to maximally support cell function—has emerged as a critical functional determinant in many biological contexts^2–20^. However, it remains unclear how aspartate levels change over time during various contexts of aspartate limitation, and how these changes are registered on downstream metabolic fates and cell function. A more nuanced understanding of these factors is important to both better understand the integration of aspartate with metabolic state and to identify opportunities to treat human diseases that interface with aspartate limitation.

## Results

### Temporally resolved measurements of aspartate levels and proliferation reveal nuanced and distinct dynamics across aspartate limitation paradigms

To enable time-resolved, nondestructive measurement of relative aspartate abundance in living cells, we employed jAspSnFR3^21^, a genetically encoded aspartate biosensor which consists of an engineered aspartate binding domain (ABD) linked to a circularly permuted green fluorescent protein (cpGFP) that is activated upon aspartate binding at the ABD (Figure 1a). Expression of jAspSnFR3 along with a nuclear-localized variant of the red fluorescent protein mRuby2 (NucRFP) produces an experimental system where relative cytosolic aspartate abundance is reported by the GFP/RFP intensity ratio and cellular proliferation rates between timepoints can be calculated using nuclei instances (Figure 1a)^21^. Measuring both variables simultaneously by live-cell imaging therefore allows us to dissect the temporal relationship between aspartate abundance and cell proliferation in unprecedented detail.

**Figure 1.**
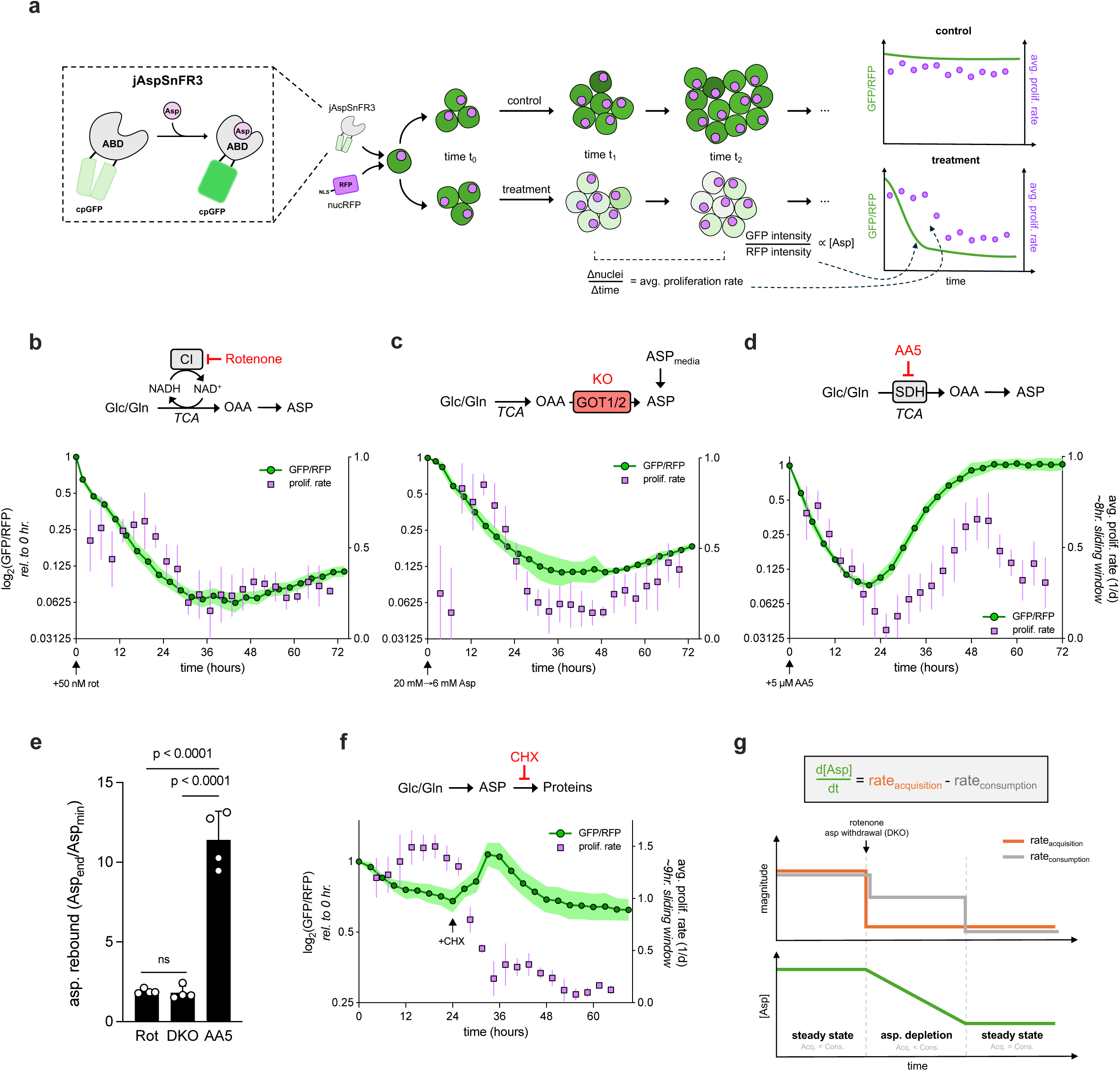
Temporally resolved measurements of aspartate levels and proliferation rate reveal distinct dynamics across aspartate limitation paradigms. **a**, schematic illustrating the experimental setup allowing simultaneous measurement of relative aspartate levels (jAspSnFR3) and proliferation rates (NucRFP) of cells in culture. GFP/RFP intensities correspond to relative aspartate levels and average proliferation rates are calculated using the equation (log_2_(nuclei_t2_/nuclei_t1_))/(t_2_-t_1_) (see Methods). **b**, relative aspartate levels (green line, left y-axis) and absolute proliferation rates (purple points, right y-axis) of 143B cells in DMEM without pyruvate treated with the CI inhibitor rotenone (50 nM), which indirectly blocks aspartate synthesis by preventing NAD^+^ regeneration by complex I (CI) (n=4). **c**, relative aspartate levels and absolute proliferation rates of GOT1/2 DKO 143B cells after switching from 20 mM to 6 mM aspartate in DMEM without pyruvate (n=4). **d**, relative aspartate levels and absolute proliferation rates of 143B cells in DMEM with 1 mM pyruvate treated with the SDH inhibitor Atpenin A5 (AA5, 5 μM), which directly blocks aspartate synthesis by inhibiting oxidative TCA cycling (n=4). **e**, comparison of the degree of aspartate rebound, calculated by dividing the GFP/RFP at 72 hours by the minimal GFP/RFP measured during an experiment, between the three aspartate limitation paradigms in b-d (n=4). **f**, relative aspartate levels and absolute proliferation rates of 143B cells treated with the translation inhibitor cycloheximide (CHX, 1 μg/mL) at 24 hours (n=4). **g**, schematic illustrating a model in which aspartate dynamics are determined by rates of aspartate acquisition and consumption, relevant to rotenone and GOT1/2 DKO experiments. Data represented as mean +/− S.D. Statistical significance determined using an ordinary one-way ANOVA with uncorrected Fisher’s LSD and a single pooled variance. ABD, aspartate binding domain; cpGFP, circularly permuted green fluorescent protein; NLS, nuclear localization signal; CI, respiratory complex I; Glc, glucose; Gln, glutamine; TCA, tricarboxylic acid cycle; OAA, oxaloacetate; ASP, aspartate; SDH, succinate dehydrogenase; DKO, GOT1/2 double knockout

First, we used this system to examine aspartate/proliferation dynamics in rapidly dividing cells under standard culture conditions. Clonal 143B osteosarcoma cells expressing jAspSnFR3/NucRFP (hereafter referred to as ‘sensor cells’) were seeded onto multi-well plates, acclimated for 24 hours, and then imaged approximately every three hours for a total of ∼72 hours in an Incucyte live cell imaging system (see Methods). To assess relative intracellular aspartate levels, total integrated GFP intensity is normalized to the total integrated RFP intensity per-well. Meanwhile, nuclei counts are used to calculate average proliferation rates in a sliding ∼8-hour window throughout the duration of the experiment and plotted alongside relative GFP/RFP intensities (Extended Data Figure 1a). In control treatments, aspartate levels remained high and relatively unchanged, consistent with the lack of any perturbation and with the notion that cellular aspartate metabolism is at homeostasis. Likewise, for the first ∼48 hours of the assay, proliferation rates remained stable at expected values for this cell line^7^, after which they become variable and slightly decrease, likely resulting from cells reaching confluency (Extended Data Figure 1b, Supplemental Video 1). Consistent with the notion that the cells are operating with finite nutrient and space constraints, repeating the experiment in DMEM supplemented with 1 mM pyruvate—a common media additive which can increase cell proliferation—leads to a more pronounced tapering of proliferation after 48 hours, corresponding to confluence, as well as a gradual decrease in aspartate abundance, potentially due to nutrient depletion (Extended Data Figure 1c-d, Supplemental Video 2)^21^. Overall, these results argue that dual expression of jAspSnFR3 and NucRFP does not significantly alter cell proliferation or metabolism, and that aspartate levels and proliferation are relatively stable in unperturbed cells pre-confluency.

Next, we leveraged this system to investigate temporal changes in aspartate levels and proliferation in several paradigms of aspartate limitation. First, we conducted a similar experiment on sensor cells cultured in DMEM without pyruvate and treated with rotenone, an electron transport chain (ETC) complex I inhibitor that blocks NAD^+^ regeneration from NADH, thereby slowing NAD^+^-dependent reactions in the TCA cycle and impairing aspartate synthesis^2–4,9^ (Figure 1b). Treatment with a dose of rotenone that robustly inhibits CI activity, but whose antiproliferative effects in 143B cells are rescuable with exogenous aspartate^2,7^ (Extended Data Figure 2a), caused a rapid decay in cellular aspartate levels, which stabilized at a lower abundance after approximately 36 hours (Figure 1b, Extended Data Figure 1e, Supplemental Video 3). In contrast, proliferation rates remained relatively stable at ∼0.5 doublings/day (roughly half that of unperturbed 143B cells) for the first 18 hours following rotenone treatment, at which point they shifted to a different, lower pseudo-steady state of ∼0.25 doublings/day for the remainder of the assay (Figure 1b). These results reveal nuanced dynamics in aspartate levels and proliferation that would otherwise be masked in single-timepoint LCMS experiments or conventional proliferation assays and indicate that cell proliferation responds to rotenone-induced aspartate limitation in two distinct phases.

Next, we sought a paradigm of aspartate limitation that doesn’t rely on pharmacological perturbation. To this end, we leveraged 143B cells lacking both glutamic-oxaloacetic transaminases 1 and 2 (GOT1 and GOT2)—which are aspartate auxotrophs^21^—and generated clonal lines of these cells expressing jAspSnFR3/NucRFP (hereafter referred to as ‘GOT1/2 DKO sensor cells’) (Extended Data Figure 2b). While GOT1/2 DKO sensor cells are maintained in 20-40 mM aspartate to maximally support cell proliferation, in agreement with previously measured parameters for non-specific aspartate uptake^16,21,22^, titrating media aspartate dose-dependently reduced their proliferation rate (Extended Data Figure 2c), outlining an experimental system in which aspartate acquisition can be modulated independent of its production. We imaged GOT1/2 DKO cells that had acclimated to DMEM containing excess (20 mM) aspartate upon switching into media with 6 mM aspartate—a concentration which restrains proliferation without overt lethality (Extended Data Figure 2c). Similar to rotenone, aspartate withdrawal in this system led to an immediate and steady decay of aspartate levels which largely stabilized by 36 hours and increased marginally by the end of the assay (Figure 1c, Supplemental Video 4). While proliferation rates were variably low for the first 6 hours, likely due to cells acclimating after the saline wash at time zero (necessary to remove residual aspartate), proliferation quickly stabilized at a moderately high rate (∼0.75 doublings/day) before shifting to a second, lower pseudo steady state of ∼0.25 doublings/day for the remainder of the assay (Figure 1c).

Overall, aspartate limitation induced by rotenone treatment or aspartate starvation in engineered auxotrophs produces similar aspartate/proliferation dynamics with three salient features: 1) proliferation defects lag changes in aspartate levels, with either paradigm showing an initial ∼12-24 hour time window in which aspartate levels significantly decrease while proliferation remains constant, 2) proliferation rates show a biphasic pattern, with a higher pseudo-steady state transitioning into a second, lower pseudo-steady state shortly before aspartate levels approach a local minimum, and 3) at the end of the assay, cells exist in a regime where both aspartate levels and proliferation rate are low relative to their initial magnitudes.

Finally, we tested a third paradigm of aspartate limitation centered on inhibition of succinate dehydrogenase (SDH), a TCA cycle enzyme implicated in several cancer types. Notably, SDH loss can also cause aspartate limitation, with metabolic features distinct from other ETC inhibitors^7^. We acclimated sensor cells in DMEM with pyruvate before treating with Atpenin A5 (AA5)—a potent and specific SDH inhibitor that directly inhibits aspartate synthesis from oxidative TCA cycling^6–8^ — and tracked the resulting aspartate/proliferation dynamics (Figure 1d). Initial aspartate and proliferation kinetics were largely concordant with the other two aspartate limitation paradigms, with a monotonic decrease in aspartate levels accompanying a slightly delayed decrease in proliferation; however, instead of leveling out at a minimum, aspartate levels surprisingly rebounded starting at ∼24 hours, reaching a GFP/RFP signal comparable to their initial values by around 48 hours (Figure 1d, Extended Data Figure 1f, Supplemental Video 5). To better understand the extent of this aspartate rebound in the three aspartate limitation paradigms, we calculated an ‘aspartate rebound’ metric by dividing the final GFP/RFP by the minimum GFP/RFP during the assay per replicate. Comparing this metric among the three paradigms reveals that the relatively modest rebounds in aspartate signal in rotenone-treated cells and aspartate-starved GOT1/2 DKO cells are dwarfed by the rebound in AA5-treated cells (Figure 1e). Proliferation dynamics upon AA5 treatment also departed from those of rotenone-treated/DKO cells, hitting a local minimum shortly after 24 hours, rebounding by ∼48 hours, and then decreasing again by the end of the assay (Figure 1d). These dynamics differ from the other two aspartate limitation paradigms both in that aspartate levels/proliferation rates rebound following an initial decay, and in that cells eventually settle into a regime where aspartate levels are partially restored, but proliferation rates remain low.

To generalize these findings beyond 143B cells, we established and analyzed the three aforementioned aspartate limitation paradigms in H1299 non-small cell lung cancer cells expressing jAspSnFR3/NucRFP (Extended Data Figure 2b)^21^. While maximal proliferation rates, aspartate uptake rates, and drug sensitivities differ in this system, aspartate/proliferation dynamics in all three paradigms were comparable to those of 143B cells (Extended Data Figure 2d-j).

All three aspartate limitation paradigms suggest that aspartate levels initially drop due to reduced aspartate acquisition (biosynthesis or uptake), while cells continue to consume their aspartate reserves as they generate biosynthetic intermediates to support proliferation. To test whether reducing aspartate consumption without impairing its acquisition would lead to the converse dynamics (i.e. an accumulation of aspartate), we treated sensor cells with the protein synthesis inhibitor cycloheximide (CHX) 24 hours after the start of an imaging assay. Indeed, CHX treatment caused a rapid decrease in proliferation rate and a corresponding spike in relative aspartate abundance (Figure 1f), which remained high but decreased towards the end of the assay, likely from feedback inhibition of aspartate synthesis. These results indicate that cellular aspartate dynamics are responsive to both aspartate acquisition and consumption and illustrate that aspartate levels and proliferation rate are not inherently coupled.

In aggregate, our experiments are consistent with a model in which aspartate dynamics in proliferating cells are determined by rates of aspartate acquisition (biosynthesis or uptake) and consumption (Figure 1g). Cells in standard conditions have matched, high levels of aspartate acquisition and aspartate consumption, leading to a stable aspartate concentration over time. Interventions that impair aspartate acquisition have an immediate but moderate effect on cell proliferation; however, there is an initial time period during which proliferation (which is inherently proportional to aspartate consumption) remains constant, leading to a depletion of aspartate pools. At least in the case of CI inhibition or aspartate starvation in GOT1/2 DKO cells, aspartate pools continue to decrease until they reach a threshold at which aspartate presumably becomes limiting for macromolecular synthesis; aspartate consumption then decreases to match production, and the cells enter a new pseudo-steady state of matched, low aspartate acquisition and consumption (Figure 1g).

While CI inhibition, GOT1/2 DKO aspartate starvation, and SDH inhibition all exert comparable long-term (i.e. 3-4 day), aspartate-dependent antiproliferative effects (Extended Data Figure 2a), our results suggest that cellular aspartate and proliferation dynamics are strikingly distinct upon SDH inhibition compared to the other two paradigms. These findings hinted at differences in aspartate acquisition or consumption following SDH inhibition that we further investigated.

### SDH inhibition impairs aspartate utilization into pyrimidine biosynthesis

To confirm the authenticity of the aspartate rebound measured by jAspSnFR3, we quantified relative whole-cell aspartate abundance using liquid chromatography-mass spectrometry (LC-MS) at different timepoints following AA5 or vehicle treatment. Consistent with biosensor measurements, aspartate levels were significantly higher at 44 hours than 24 hours following AA5 treatment (Figure 2a). We next sought to rule out the possibility that the aspartate rebound was caused by degradation of AA5 and disinhibition of SDH during the imaging assays. To this end, we quantified AA5 itself as well as the succinate/fumarate ratio—a metabolic surrogate for SDH activity^23^—at several timepoints following AA5 treatment. AA5 was undetectable in vehicle-treated cells, and AA5 abundances remained high between 10-66 hours post-treatment (Extended Data Figure 3a), arguing against meaningful drug degradation. Meanwhile, AA5 treatment caused a >1,000-fold increase in the succinate/fumarate ratio compared to vehicle-treated controls at all timepoints, indicating durable SDH inhibition over the entire assay (Extended Data Figure 3b).

**Figure 2.**
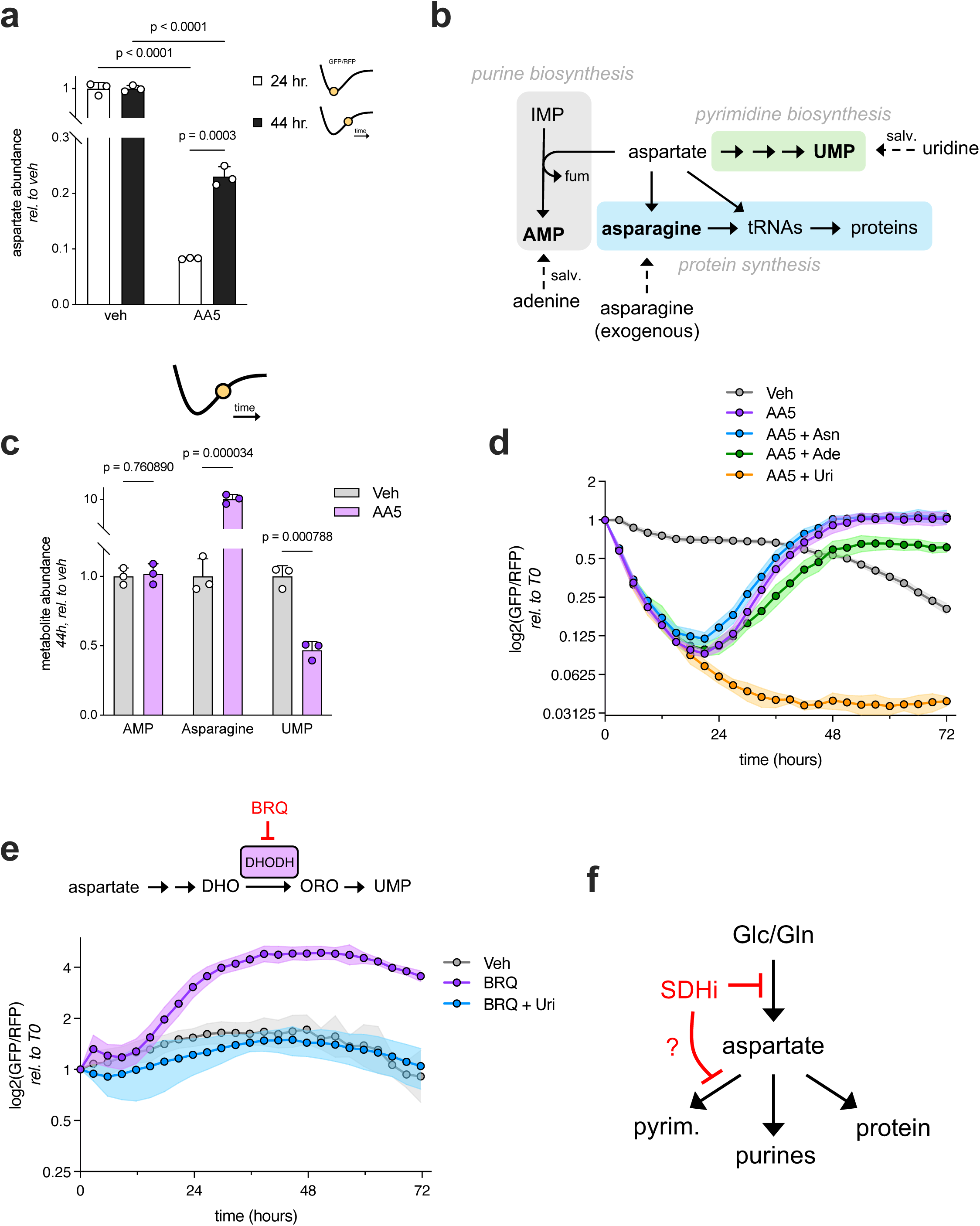
SDH inhibition impairs aspartate utilization into pyrimidine biosynthesis. **a**, aspartate levels measured using LC-MS of 143B cells at 24– and 44-hours post-treatment with 5 μM AA5, each relative to vehicle control (DMSO) (n=3). Cartoons illustrate the approximate location of each timepoint on the prototypical GFP/RFP rebound curve in AA5-treated cells **b**, schematic illustrating aspartate fates in our system, including metabolites which can be salvaged (salv.) into each fate. **c**, relative abundances of AMP, Asparagine, and UMP measured using LC-MS of 143B cells at 44-hours post-treatment with 5 μM AA5 or vehicle control (DMSO) (n=3). The cartoon indicates the position of this timepoint on the GFP/RFP rebound curve in AA5-treated cells. **d**, relative aspartate levels of 143B sensor cells treated with vehicle control (DMSO), 5 μM AA5 alone, or 5 μM AA5 supplemented with 500 μM asparagine (Asn), 100 μM adenine (Ade), or 200 μM uridine (Uri) (n=4). **e**, relative aspartate levels of 143B sensor cells treated with vehicle control (DMSO), 2 μM of the DHODH inhibitor brequinar (BRQ), or 2 μM BRQ and 200 μM uridine (BRQ + Uri) (n=4). **f**, schematic illustrating the hypothesis that SDH inhibition simultaneously impairs aspartate synthesis and consumption into pyrimidine biosynthesis. Unless otherwise noted, experiments were conducted in DMEM with 1 mM pyruvate. Data represented as mean +/− S.D. Statistical significance determined using an ordinary two-way ANOVA with uncorrected Fisher’s LSD and a single pooled variance (panel a) or multiple unpaired t-tests (panel c). IMP, inosine monophosphate; AMP, adenosine monophosphate; UMP, uridine monophosphate; DHO, dihydroorotate; ORO, orotate; fum, fumarate.

Under the model of aspartate dynamics introduced in the previous section (Figure 1g), the aspartate rebound ought to arise from increased aspartate acquisition, decreased aspartate consumption, or a combination of the two. The short timescale in which the rebound occurs, lack of aspartate in the culture media, and sustained SDH inhibition by AA5 during the assay (Extended Data Figure 3a, b) argue against an increase in aspartate uptake or synthesis via the canonical route. Instead, we turned our attention to aspartate consumption. Aspartate is consumed for the biosynthesis of proteins (via charging aspartyl-tRNAs), asparagine, and both purine and pyrimidine nucleotides (Figure 2b). To determine whether aspartate consumption into protein synthesis may be affected upon SDH inhibition, we quantified the charge of aspartyl– and asparaginyl-tRNAs in 143B cells treated with AA5 or vehicle using tRNA-seq^24^. This analysis revealed sustained high charging of all relevant tRNA species in AA5-treated cells, arguing against aspartate limitation into protein synthesis as a driver of the aspartate rebound (Extended Data Figure 3c).

To evaluate the consumption of aspartate in SDH-inhibited cells into asparagine, purine, and pyrimidine biosynthesis, respectively, we quantified relative abundances of asparagine, AMP, and UMP at 44-hours post AA5 treatment (when aspartate levels are ∼mid-rebound). While AMP levels were unchanged and asparagine levels were significantly higher, UMP levels were depleted in AA5-treated cells compared to vehicle controls, suggesting that aspartate consumption into pyrimidines is reduced during the aspartate rebound (Figure 2c). For a closer look at pyrimidine biosynthesis, we quantified the relative abundances of the pyrimidine intermediates carbamoyl-aspartate, dihydroorotate, orotate, and UMP at 24, 44, and 66 hours following AA5 treatment in 143B cells using LC-MS. Strikingly, all four were depleted in AA5-treated cells relative to vehicle-treated controls at each timepoint (Extended Data Figure 3d), further suggesting impaired aspartate consumption into *de novo* pyrimidine synthesis upon SDH inhibition. Interestingly, levels of these metabolites increased towards the end of the experiment, suggesting that pyrimidine synthesis partially recovers over time.

Next, we sought to exogenously fulfill each individual aspartate fate and determine the effects on aspartate dynamics following SDH inhibition. Since each fate has discrete uptake and incorporation characteristics, we first investigated the nutrient conditions necessary to bypass the demand for aspartate consumption for the synthesis of asparagine, pyrimidines, and purines. To do so, we used isotope tracing strategies in 143B and H1299 cells to label endogenously produced aspartate fates and then determined the metabolite supplementation conditions that could displace the labeled species, indicating that *de novo* synthesis from aspartate was no longer necessary. As expected, treatment with unlabeled asparagine suppressed *de novo* asparagine synthesis and treatment with unlabeled uridine robustly suppressed the contribution of *de novo* pyrimidine synthesis to the UTP pool (Extended Data Figure 4a, b). Although aspartate can serve as a nitrogen donor for arginine biosynthesis, many cancer cell lines suppress arginine synthesis and instead import arginine from culture media^19,25^. Indeed, label incorporation was undetectable from glutamine into arginine (Extended Data Figure 4c). Among purine nucleobase treatments, adenine was sufficient to meet all purine demands, bypassing the aspartate consumption step specific to adenylate nucleotide synthesis and supporting guanylate nucleotide production, presumably through deamination to generate IMP, and bypassing the earlier aspartate consumption step common to all *de novo* synthesized purines (Extended Data Figure 4d). Altogether, this investigation identified concentrations of exogenous adenine, uridine, and asparagine that can efficiently fulfill the non-protein metabolic fates of aspartate in this system (Figure 2b).

We then conducted a 72-hour imaging assay in which sensor cells were treated with AA5 and either asparagine, adenine, or uridine to determine the effects on aspartate dynamics. While adenine and asparagine did not substantially affect the aspartate rebound, uridine supplementation completely abolished it, resulting in monotonic aspartate depletion over the course of the assay (Figure 2d). Using LC-MS, we confirmed that uridine supplementation significantly reduced aspartate levels in AA5-treated cells and restored UTP levels (Extended Data Figure 3e, f), corroborating the sensor results and indicating that pyrimidine deficiency is necessary for the aspartate rebound effect upon SDH inhibition.

Finally, we sought to determine whether inhibition of pyrimidine synthesis is sufficient to increase aspartate levels. To this end, we treated sensor cells with brequinar (BRQ), a specific inhibitor of the DHODH step of pyrimidine synthesis^26^ and measured the resulting aspartate dynamics. Strikingly, BRQ treatment alone caused a spike in aspartate levels with similar temporal kinetics as the AA5-mediated rebound and was negated by uridine co-treatment, confirming the on-target effect of BRQ (Figure 2e). Altogether, these results argue that aspartate levels rebound following SDH inhibition due to a specific impairment of pyrimidine nucleotide biosynthesis. The fact that pyrimidine biosynthesis intermediates remain depleted from 24-44 hours post AA5-treament (Extended Data Figure 3d) despite a significant increase in aspartate levels during this time period (Figure 2a) suggests that SDH inhibition causes a secondary metabolic effect that impairs *de novo* pyrimidine biosynthesis beyond simply limiting aspartate availability (Figure 2f).

### Distributed and hierarchical metabolic growth limitations upon SDH inhibition

The fact that salvageable aspartate fates differed in preventing the aspartate rebound posed the question of how their supplementation impacts cell proliferation upon SDH inhibition. Interestingly, while adenine and asparagine did not substantially affect proliferation dynamics following AA5 co-treatment (Extended Data Figure 5b, c)—consistent with their lack of impact on aspartate dynamics (Figure 2d)—uridine prevented the decrease in proliferation rate during the initial 24 hours following AA5 treatment, after which proliferation rates monotonically decreased instead of rebounding as observed in the other conditions (Extended Data Figure 5a). These data suggest that uridine initially solves the proximal proliferation defect from SDH inhibition, but that cells then run into a secondary, aspartate-related metabolic limitation as aspartate levels are further consumed for proliferation. Indeed, LC-MS at 32 hours following AA5/uridine co-treatment revealed that AMP (but not asparagine) was significantly depleted in AA5/uridine co-treated cells relative to controls (Extended Data Figure 5d). IMP, the metabolite immediately upstream of the aspartate-dependent step in purine synthesis (Figure 2b), was also drastically accumulated, consistent with a purine deficiency due to reduced aspartate levels impairing IMP to AMP conversion, as has been found in other settings of aspartate limitation^2,16^ (Extended Data Figure 3e).

To verify that these cells were not also deficient in the proteogenic aspartate fates at this time, we measured relative protein synthesis rates using a puromycin incorporation assay 24 hours following treatment with vehicle, AA5, or AA5/uridine. While both AA5-containing treatments had reduced puromycin incorporation relative to vehicle-treated cells, there was no significant difference between AA5 and AA5/uridine treated conditions (Extended Data Figure 5e, f), indicating that uridine treatment does not further impair protein synthesis in AA5-treated cells at this timepoint.

To functionally test whether purine deficiency is responsible for the proliferation decrease upon AA5/uridine co-treatment, we measured aspartate/proliferation dynamics in cells treated with AA5 and both uridine and adenine. Uridine/adenine co-supplementation further improved proliferation to ∼1 doubling/day for 24 hours following AA5 treatment, after which proliferation shifted to a lower but relatively stable ∼0.5 doublings/day for the remainder of the assay (Extended Data Figure 5g), consistent with uridine supplementation precipitating a secondary purine deficiency in SDH-impaired cells. Notably, GFP/RFP levels still monotonically decreased throughout the assay, suggesting that continued aspartate consumption into its other fates (asparagine/protein synthesis) still outpaced acquisition in this context.

Finally, to test whether the proliferation shift in AA5/uridine/adenine treated cells may reflect an emergent deficiency in asparagine synthesis upon pyrimidine/purine rescue, we measured aspartate/proliferation dynamics after additionally supplementing with asparagine. Consistent with this hypothesis, proliferation rates in AA5/uridine/adenine/asparagine treated cells were sustained at around 0.8 doublings/day throughout the entire assay, while aspartate levels still decayed but plateaued at relatively higher levels (Extended Data Figure 5h). Overall, these results reveal that the metabolic growth limitations in cells upon SDH inhibition are 1.) distributed, with no single aspartate fate able to provide a sustained proliferation benefit in the absence of the other fates, but also 2.) hierarchical, with pyrimidine deficiencies superseding purine deficiencies, which supersede deficiencies in the proteogenic aspartate fates. This is consistent with a model whereby aspartate fates are all required for sustained cellular proliferation despite exhibiting different ‘aspartate thresholds’ at which they become impaired. Rescuing a single fate permits continued aspartate consumption until aspartate levels become limiting for the next fate in the hierarchy (Extended Data Figure 5i).

### Succinate inhibits mammalian ATCase

In SDH-impaired cells, pyrimidine precursors are depleted and remain so even as aspartate levels recover, suggesting that SDH inhibition alters metabolism in a way that specifically disfavors pyrimidine synthesis beyond simply limiting aspartate availability. A key metabolic distinction between SDH inhibition and other causes of aspartate limitation is the accumulation of succinate, the substrate of SDH. In addition to its role as a TCA cycle intermediate, succinate is an ‘oncometabolite’ known to have multiple biochemical effects, including competitively inhibiting various alpha ketoglutarate-dependent dioxygenases involved in oxygen sensing and DNA/histone demethylation^27^. To rigorously determine the effects of SDH inhibition on succinate and aspartate in our system, we used quantitative LC-MS to estimate whole-cell concentrations of these metabolites in 143B cells treated with AA5 or vehicle control, combining multiple treatment durations. We found that vehicle-treated 143B cells maintain median succinate levels around 200 μM and aspartate levels around 1.7 mM, consistent with other measurements of unperturbed mammalian cells^21,28^. SDH inhibition increased succinate levels by approximately 70-fold, reaching a median concentration of around 14 mM, which is comparable to what has been measured in SDH-deficient tumors and cell lines^29–32^ (Figure 3a). Meanwhile, SDH inhibition decreased aspartate concentrations by roughly 8-fold—with some variability depending on timepoint—to a median concentration of approximately 200 µM (Figure 3b). These results confirm that SDH inhibition in our system both depletes aspartate and dramatically increases cellular succinate concentrations, motivating us to search for mechanistic links between succinate and pyrimidine biosynthesis.

**Figure 3.**
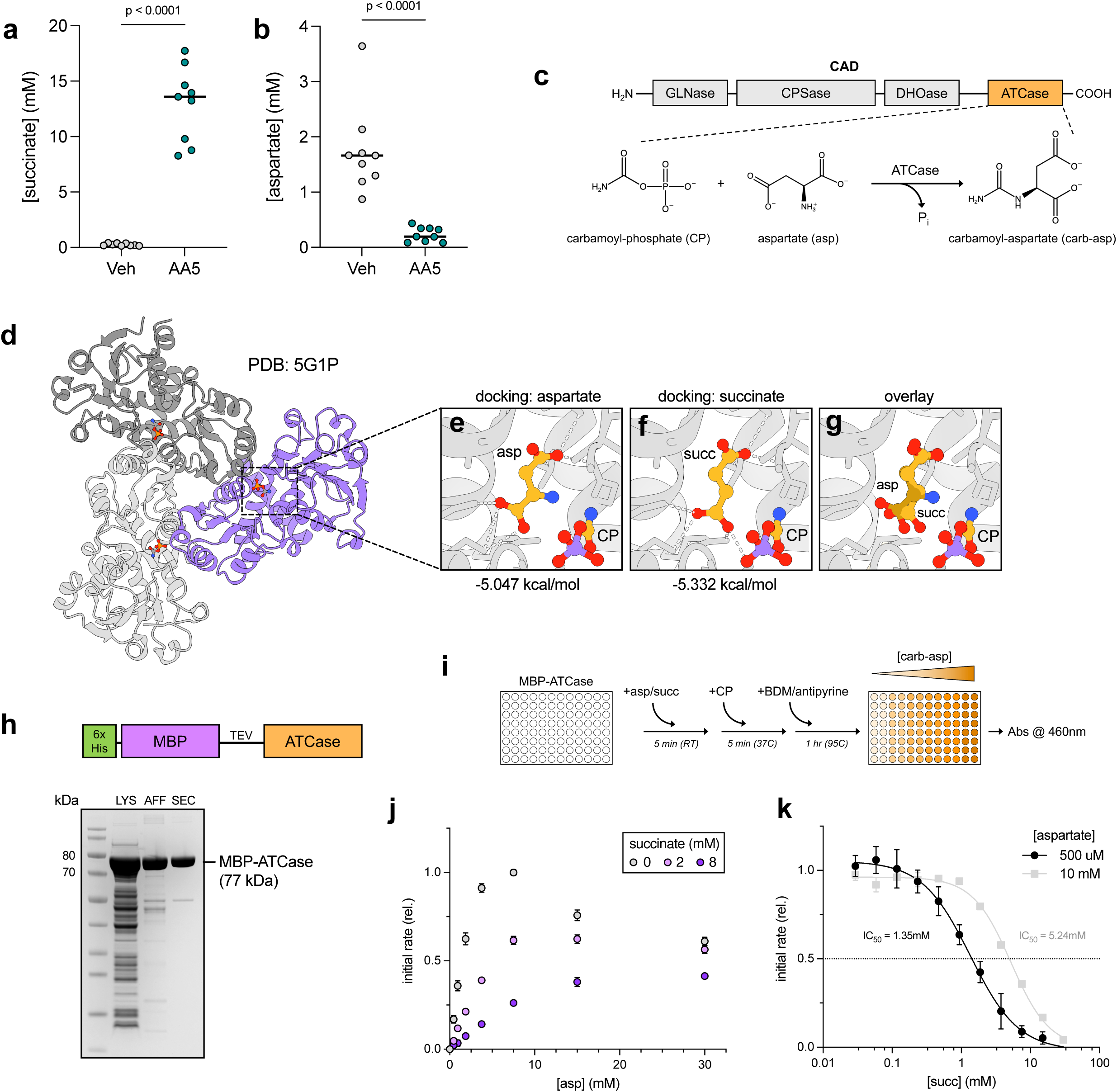
Succinate inhibits mammalian ATCase. **a-b**, whole-cell aspartate (b) and succinate (c) concentrations in 143B cells measured using LC-MS at multiple timepoints following treatment with vehicle or 5 μM Atpenin A5 (AA5) (n=9, multiple timepoints from several different LC-MS experiments). **c**, a schematic of the mammalian CAD protein with the C-terminal ATCase domain highlighted, and the reaction catalyzed by ATCase. **d**, a previously-solved crystal structure of the mammalian ATCase trimer (PDB: 5G1P) bound to carbamoyl-phosphate (CP), colored by monomer, which was used for molecular docking analyses. **e-g**, predicted binding poses and affinities of aspartate (c), succinate (d), and aspartate/succinate (e) in the substrate binding pocket of CP-bound mammalian ATCase. Ligands are colored by atom: yellows = carbon, red = oxygen, blue = nitrogen, purple = phosphorus. **h**, a schematic of the 6xHis-tagged MBP-ATCase construct used in subsequent panels, and a Coomassie-stained protein gel demonstrating expression and purification of MBP-ATCase. LYS = cleared bacterial lysate; AFF = elute following affinity purification; SEC = elute following size-exclusion chromatography. **i**, schematic illustrating the absorbance-based ATCase activity assay which measures accumulation of the reaction product, carbamoyl-aspartate (see Methods). **j**, representative activity assay depicting initial enzymatic rates of purified MBP-ATCase incubated with 10 mM CP and the indicated concentrations of aspartate (asp) and succinate. Rates are normalized to the 0 mM succinate, 7.5 mM aspartate condition (n=3 technical replicates). **k**, representative activity assay depicting initial enzymatic rates of purified MBP-ATCase incubated with 10 mM CP and the indicated concentrations of succinate (succ) at either 500 μM or 10 mM aspartate (n=3 technical replicates). Rates are normalized to the 0 mM succinate conditions for each respective aspartate concentration and fit to a sigmoidal curve using Prism. Data represented as mean +/− S.D. Statistical significance determined using unpaired t-tests. MBP, maltose binding protein; TEV, Tobacco etch virus protease site; BDM, butanedione monoxime.

SDH inhibition depletes carbamoyl-aspartate and downstream pyrimidine intermediates (Extended Data Figure 3d), suggesting that impairment occurs at the first step of pyrimidine synthesis—the formation of carbamoyl-aspartate from aspartate and carbamoyl-phosphate— which is catalyzed by aspartate transcarbamylase (ATCase) (Figure 3c). Interestingly, several classic enzymological studies noted that succinate can competitively inhibit *Escherichia coli* ATCase in vitro with a K_i_ in the range of ∼0.4-20 mM, depending on pH^33–35^. To our knowledge, succinate inhibition of mammalian ATCase has not been described, nor has this interaction been observed in living cells. Nonetheless, as bacterial and human ATCase share significant sequence and structural similarity, including at the active site^36^ (Extended Data Figure 6a), we reasoned that succinate-mediated inhibition of ATCase could explain the effects of SDH impairment on pyrimidine biosynthesis.

Despite sharing a catalytic fold, bacterial and human ATCase are non-identical and differ in some important respects. Bacterial ATCase is a standalone enzyme which forms catalytic homotrimers flanked by regulatory subunits, which dimerize to form dodecamers, while human ATCase comprises the C-terminal domain of a mega-enzyme called CAD (carbamoyl-phosphate synthetase 2, aspartate transcarbamylase, and dihydroorotase) that combines the catalytic activities of the first three steps in pyrimidine biosynthesis and likely trimerizes via the ATCase domain^37^ (Figure 3c). To evaluate if succinate could bind the ATCase substrate pocket in a manner similar to aspartate, we performed molecular docking analyses using a published x-ray crystal structure of the ATCase domain of human CAD^36^. For this analysis, we chose the structure of CP-bound human ATCase (Figure 3d), since CP binds ATCase before aspartate and prepares the binding pocket to accept aspartate^36,38^. Since we could not find a structure of human ATCase bound to both CP and aspartate, we first docked aspartate into the substrate pocket of CP-bound ATCase. This analysis revealed several top-scoring poses which orient the bound aspartate with its α-amino group proximal to the carbonyl of CP, consistent with the proposed ATCase reaction mechanism^38^ (Figure 3e). Repeating this docking analysis with succinate returned two top-scoring poses in which succinate occupies nearly the same pose as aspartate, minus the α-amino group (Figure 3f-g, Extended Data Figure 6b). Notably, the predicted binding affinities for succinate and aspartate are similar (–5.332 and –5.047 kcal/mol, respectively), supporting the hypothesis that succinate acts as a substrate-analog inhibitor of human ATCase.

To test this hypothesis experimentally, we cloned, recombinantly expressed, and purified the ATCase domain of human CAD fused N-terminally to a 6xHis-tagged maltose binding protein (MBP) (Figure 3h). Size exclusion chromatography of affinity-purified MBP-ATCase revealed a single major peak at the expected mass of the trimer (∼241 kDa) (Extended Data Figure 6c), confirming that the MBP tag did not prevent oligomerization. Using an adapted plate-based ATCase activity assay^39^ which relies on a colorimetric readout of carbamoyl-aspartate formation (Figure 3i, Extended Data Figure 6d), we measured relative initial enzymatic rates of MBP-ATCase in excess aspartate and a titration of CP. This showed saturation kinetics that were consistent with a previous study^36^. Furthermore, initial rates were dose-dependently decreased with nanomolar doses of the classical ATCase inhibitor, PALA (N-phosphonacetyl-L-aspartate)^40^, as expected (Extended Data Figure 6e). Next, we measured initial enzymatic rates of MBP-ATCase in excess CP, a titration of aspartate, and three concentrations of succinate (0, 2, and 8 mM). This analysis revealed a potent and dose-dependent inhibition of enzyme activity by succinate (Figure 3j).

The fact that human ATCase exhibits positive cooperativity with regards to aspartate and that high aspartate concentrations paradoxically inhibit human ATCase activity^36^ (Figure 3j) prevents a straightforward modeling of our enzyme kinetic data and assigning of a single K_i_ value for succinate. Nevertheless, fitting the aforementioned activity data to sigmoidal curves suggests that succinate increases K_1/2_ and decreases V_max_, consistent with mixed inhibition (Extended Data Figure 6f); meanwhile, succinate inhibits ATCase non-competitively with regards to CP (Extended Data Figure 6g). To better delineate the relationship between enzyme activity, succinate, and aspartate, we performed an activity assay in excess CP and fixed the aspartate concentration at two values: 500 μM, slightly above the median whole-cell aspartate concentration we measured in SDH-inhibited cells (Figure 3b), and 10 mM, representing a supraphysiological aspartate concentration. Titrating succinate in these conditions revealed a sigmoidal dependence between succinate concentration and enzyme activity, with IC_50_ values for succinate of 1.35 mM and 5.24 mM at 500 μM and 10 mM aspartate, respectively (Figure 3k). Finally, we repeated this experiment with fumarate, which differs from succinate only by a central double bond, and saw no appreciable inhibition of ATCase at fumarate levels up to 30 mM (Extended Data Figure 6h). Thus, succinate specifically inhibits human ATCase in a manner that depends on aspartate concentration, with IC_50_ values that are well below the median whole-cell succinate concentrations in SDH-inhibited cells.

One mechanism by which succinate could inhibit human ATCase is by disrupting its trimerization, which is essential for catalysis since adjacent monomers contribute to substrate pocket formation^41^ (Figure 3d). To address this possibility, we used mass photometry (MP) to query the particle size distribution of purified MBP-ATCase in solution with excess CP and sub-saturating concentrations of aspartate, with or without 2 mM succinate. Under both substrate conditions, MP revealed bimodal particle size distributions consistent with an equilibrium between monomeric and trimeric MBP-ATCase. The addition of succinate did not substantially change this equilibrium—and potentially even slightly favored trimer formation (Extended Data Figure 6i)—arguing that succinate does not inhibit human ATCase by disrupting oligomerization. Overall, our *in vitro* results reveal that succinate can inhibit human ATCase in a manner which is at least partially competitive with aspartate and argue strongly for succinate accumulation as the driver of pyrimidine biosynthesis impairment in SDH-inhibited cells.

### Metabolic control of aspartate and succinate abundance defines ATCase activity in cells

We previously reported that SDH-inhibited cells benefit from co-inhibition of ETC complex I (CI), which improves cell proliferation by decreasing mitochondrial NAD^+^/NADH to promote alternative aspartate synthesis^7^. CI inhibition is also expected to decrease succinate levels by slowing the activity of the NAD^+^-dependent alpha-ketoglutarate dehydrogenase (αKGDH) enzyme^7,42^, which may benefit SDH-impaired cells by lowering succinate to disinhibit ATCase (Figure 4a). First, we confirmed that the CI inhibitor rotenone significantly decreases succinate levels in AA5-treated cells, an effect which could be partially reversed by succinate supplementation (Figure 4b). Next, we examined carbamoyl-aspartate abundance in these conditions as a metabolic indicator of ATCase activity; indeed, CI inhibition significantly increased carb-asp levels at 48 hours post-treatment, and succinate supplementation largely blunted this effect (Figure 4c). Aspartate levels were slightly lower in SDH/CI impaired cells at this timepoint and increase upon treatment with exogenous succinate (Extended Data Figure 7a), further demonstrating a metabolic interaction between impaired pyrimidine synthesis and aspartate levels. We also confirmed these effects were mostly present at 24 hours post AA5/rotenone co-treatment (Extended Data Figure 7b-d). Evaluating these treatments in 143B sensor cells, we found that rotenone co-treatment delays and suppresses the AA5-induced aspartate rebound, consistent with rotenone partially restoring pyrimidine synthesis, and that succinate addition reactivates the rebound, while uridine supplementation abolishes it completely (Extended Data Figure 7e). To further validate that rotenone restores pyrimidine synthesis to SDH-inhibited cells, we traced U-^13^C glutamine into pyrimidine synthesis intermediates at 32 hours in cells treated with vehicle, AA5, or AA5/rotenone (Extended Data Figure 7f, g). As expected, levels of most pyrimidine intermediates were significantly higher in AA5/rotenone treated cells, with isotopolog distributions consistent with increased biosynthesis from aspartate (Extended Data Figure 7g). Overall, these results indicate that CI suppression disinhibits ATCase in SDH-impaired cells by reducing succinate levels and further supports a model whereby cellular ATCase activity is influenced by succinate abundance.

**Figure 4.**
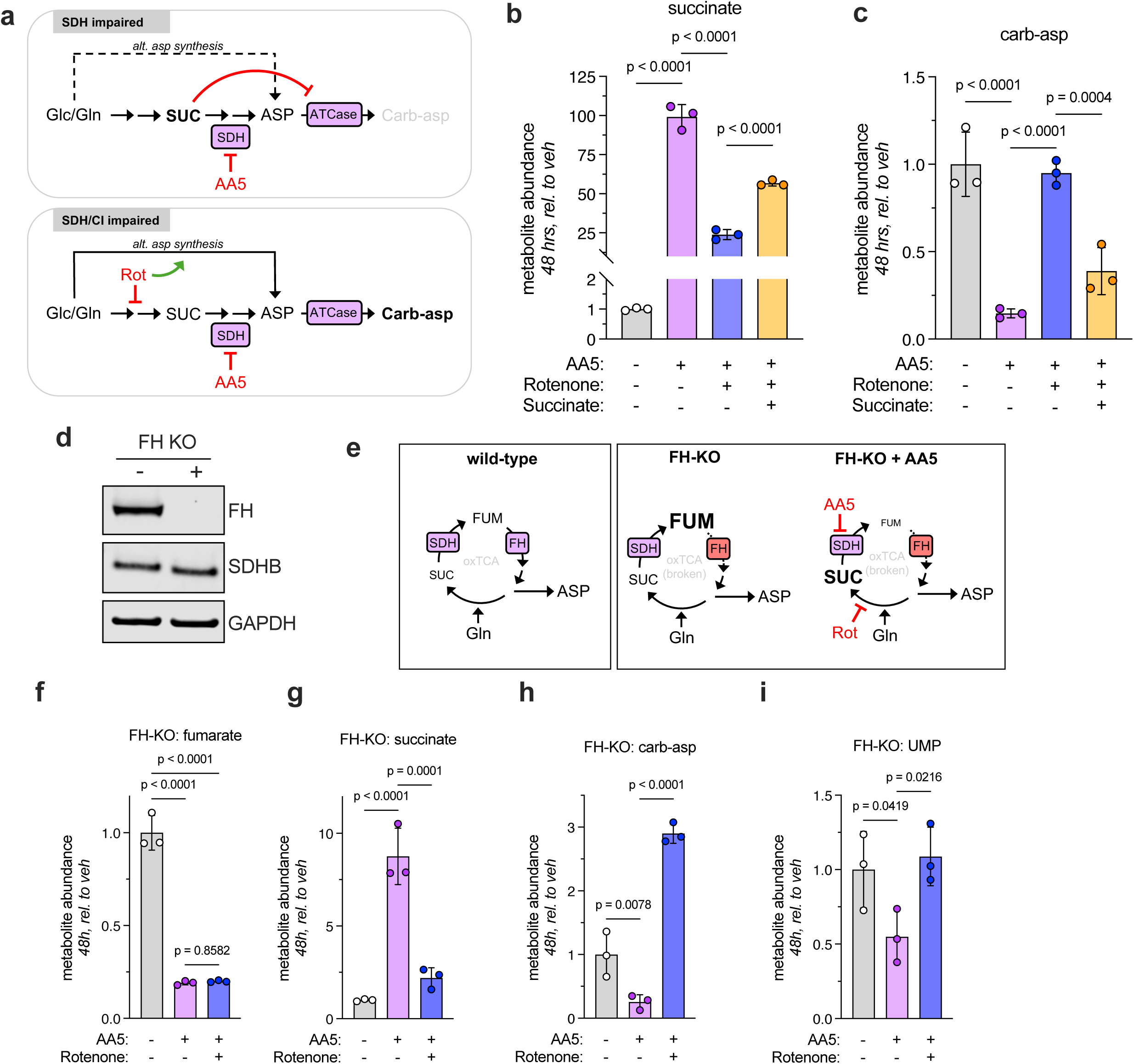
Metabolic control of aspartate and succinate abundance defines ATCase activity in cells. **a**, schematic illustrating the proposed effects of AA5 and rotenone (Rot) on aspartate synthesis and ATCase activity in cells. **b**, relative succinate abundances measured by LC-MS on 143B cells 48 hours after treatment with vehicle control or the indicated combinations of 5 μM AA5, 50 nM rotenone, and 10 mM succinate. **c**, relative carbamoyl-aspartate (carb-asp) abundances measured by LC-MS on 143B cells 48 hours after treatment with vehicle control or the indicated combinations of 5 μM AA5, 50 nM rotenone, and 10 mM succinate. **d**, Western blot showing levels of fumarate hydratase (FH), SDHB, and GAPDH loading control in 143B FH KO cells and parental 143B cells. **e**, schematic illustrating the TCA cycle status of FH-KO cells compared to wild-type parental cells and describing the effects of AA5/rotenone treatment in FH-KO cells. **f-i**, relative fumarate (f), succinate (g), carbamoyl-aspartate (h), and UMP (i) abundances measured by LC-MS on 143B FH-KO cells 48 hours after treatment with vehicle control or the indicated combinations of 5 μM AA5 and 50 nM rotenone. n=3 biological replicates for all panels, data represented as mean +/− S.D. Unless otherwise noted, experiments were conducted in DMEM with 1 mM pyruvate. Statistical significance determined using an ordinary two-way ANOVA with uncorrected Fisher’s LSD and a single pooled variance. SUC, succinate; FUM, fumarate; ASP, aspartate; Gln, glutamine; oxTCA, oxidative tricarboxylic acid cycle; UMP, uridine monophosphate.

For an orthogonal system in which we could modulate cellular succinate abundance and probe effects on ATCase activity, we used CRISPR/Cas9 to generate monoclonal 143B cells deficient in fumarate hydratase (FH), the TCA cycle enzyme immediately downstream of SDH (Figure 4d). FH-KO cells are similar to SDH-KO cells in that they have no oxidative TCA cycling; however, they differ in their preferential accumulation of fumarate, which does not inhibit ATCase *in vitro* (Extended Data Figure 6h), to a greater degree than succinate (Figure 4e)^43^. We found that treating FH-KO cells with AA5 decreased fumarate levels by nearly ten-fold and similarly increased succinate levels (Figure 4f, g), effectively transforming their predominant ‘fumarate accumulation’ phenotype into a ‘succinate accumulation’ phenotype (Figure 4e). Strikingly, AA5 treatment in FH-KO cells increased aspartate and decreased carbamoyl-aspartate levels (Extended Data Figure 7h, Figure 4h), resulting in a corresponding UMP deficiency (Figure 4i) but not deficiencies in AMP or asparagine (Extended Data Figure 7i). Rotenone co-treatment rescued these effects, consistent with their dependence on succinate accumulation (Figure 4g-i, Extended Data Figure 7h, i).

Finally, our biosensor and *in vitro* data demonstrate that succinate serves at least partially as a competitive inhibitor of ATCase, suggesting that increasing aspartate concentrations should be sufficient to re-establish ATCase activity in SDH-impaired cells. Consistent with this idea, exogenous aspartate supplementation rescued the decreases in aspartate and carbamoyl-aspartate following AA5 treatment without affecting succinate abundance (Extended Data Figure 7j-l). Altogether, these results further pinpoint aspartate and succinate as important metabolic determinants governing ATCase activity and pyrimidine biosynthesis in living cells. They also argue that pyrimidine synthesis defects are not a generalized consequence of oxidative TCA cycle dysfunction, but rather a specific consequence of the succinate accumulation that is characteristic of SDH impairment^29^.

### SDH inhibition causes replication stress by impairing pyrimidine synthesis

Imbalanced nucleotide availability can cause replication stress, a physiological state characterized by DNA replication stalling and DNA damage that can lead to impaired proliferation or cell death if not mitigated^44–47^. Since a prolonged S phase is a typical marker of replication stress^44,48^, we used propidium iodide staining and flow cytometry to characterize the cell cycles of unsynchronized 143B cells at several timepoints following treatment with AA5 or vehicle control (Figure 5a). While vehicle-treated cells showed roughly equivalent proportions of cells in G1, S, and G2 phases throughout the experiment (Figure 5b), AA5-treated cells saw a dramatic increase in the proportion cells in S phase from approximately 18 to 36 hours post-treatment (Figure 5c). Interestingly, the proportion of S phase cells then progressively decreased by 72 hours post-treatment (Figure 5c), a timeframe characterized by both aspartate and proliferation rate rebounds (Figure 1d). This S phase accumulation phenotype was rescued by uridine co-treatment (Figure 5d), indicating that it results from a pyrimidine deficiency. Indeed, inhibiting pyrimidine synthesis using BRQ also caused a similar S phase accumulation phenotype that was rescuable by uridine (Extended Data Figure 8a). In contrast to the S phase accumulation of AA5-treated cells—which begin to partially resolve around 48 hours post-treatment—BRQ-induced S phase accumulation largely persisted up to 72 hours, highlighting the distinction between BRQ’s ‘full block’ of pyrimidine synthesis at DHODH versus a competitive inhibition of pyrimidine synthesis at ATCase by succinate, which may be overcome with sufficient aspartate accumulation. We note that this ‘synchronize and release’ effect following SDH inhibition likely underlies the cell proliferation rebound observed in AA5-treated cells (Figure 1d).

**Figure 5.**
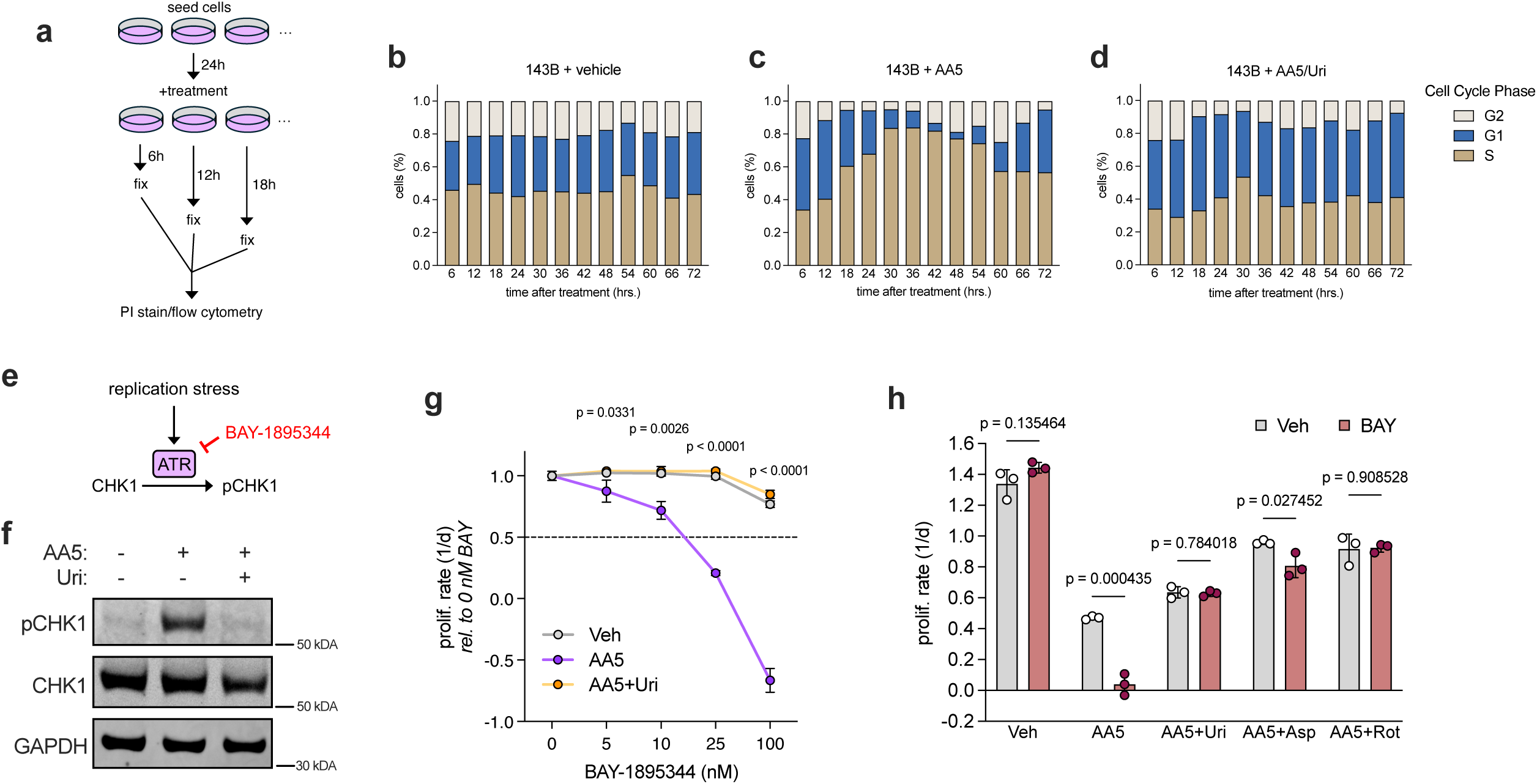
SDH inhibition causes replication stress by inhibiting pyrimidine biosynthesis. **a**, schematic illustrating the experimental setup for cell cycle analysis of 143B cells treated with AA5 for various time intervals. **b-d**, representative cell cycle experiment showing the proportion of cells in each cell cycle phase in 143B cells treated with vehicle control (b), 5 μM AA5 (c), or 5 μM AA5 and 200 μM uridine (d) at the indicated timepoints post-treatment (n=1). **e**, schematic illustrating ATR’s function in sensing replication stress and phosphorylating CHK1 in response. The ATR inhibitor BAY-1895344 (BAY) is indicated. **f**, representative Western blot demonstrating levels of phosphorylated CHK1 (pCHK1), total CHK1, and GAPDH loading control in 143B cells treated with vehicle control, 5 μM AA5, or 5 μM AA5 and 200 μM uridine for 24 hours. **g**, proliferation rates (normalized to 0 nM BAY in each respective condition and measured using a conventional, 72-hour endpoint proliferation assay) of 143B cells treated with vehicle control, 5 μM AA5, or 5 μM AA5 and 200 μM uridine and the indicated doses of BAY (n=3). **h**, absolute proliferation rates (measured using a conventional, 72-hour endpoint proliferation assay) of 143B cells treated with vehicle control or 20 nM BAY and the indicated combinations of vehicle control, 5 μM AA5, 200 μM uridine (Uri), 20 mM aspartate (Asp), and 50 nM rotenone (Rot) (n=3). Unless otherwise noted, experiments were conducted in DMEM with 1 mM pyruvate. Data represented as mean +/− S.D. Statistical significance determined using an ordinary two-way ANOVA with uncorrected Fisher’s LSD and a single pooled variance (panel g) or multiple unpaired t-tests (panel h). In panel g, p-values denote the results of statistical testing comparing Veh and AA5 conditions at each dose of BAY.

In cancer cells, cell cycling defects caused by nucleotide imbalance-mediated replication stress can occur without commensurate decreases in growth signaling, resulting in an increase in cell size^47^. Consistent with these studies, we noted that AA5 treatment significantly increases cell size—a phenotype which can be partially rescued with rotenone, re-established with succinate addition, and completely rescued by uridine supplementation (Extended Data Figure 8b). FH-KO cells are similarly sized to their parental cells but swell significantly upon AA5 treatment. This enlargement could be rescued with aspartate or uridine (Extended Data Figure 8b), supporting increased cell size as a parameter that is phenotypically linked to pyrimidine synthesis and replication stress in these contexts.

To directly evaluate replication stress signaling in response to SDH inhibition, we measured phosphorylation of the canonical replication stress-associated checkpoint kinases 1 and 2 (CHK1/2)^49^ (Figure 5e). As a positive control, we treated 143B cells with excess adenine, which causes replication stress secondary to a purine nucleotide imbalance, leading to CHK1 and subsequent CHK2 phosphorylation over time, as previously reported^47^ (Extended Data Figure 8c). Notably, AA5 treatment induced CHK1 phosphorylation as early as 24 hours post-treatment (Figure 5f, Extended Data Figure 8c), and pCHK1 levels were diminished by 72 hours, at which point AA5-treated cells begin to overcome S-phase arrest (Extended Data Figure 8c, Figure 5c). AA5-induced CHK phosphorylation could also be substantially rescued by uridine, aspartate, or rotenone—metabolically distinct treatments which all converge on alleviating pyrimidine deficiency (Figure 5f, Extended Data Figure 8c, d).

ATR signaling is essential during replication stress to prevent a catastrophic loss of DNA integrity and subsequent cell death, quiescence, or senescence^44,49,50^. Thus, we tested whether SDH impairment sensitized cells to the ATR inhibitors BAY-1895344^51^ (BAY) or VE-821^52^, which we verified to block CHK1 phosphorylation (Extended Data Figure 8e, f). Indeed, while up to 100 nM BAY had little effect on 143B cell proliferation, AA5 treatment dramatically sensitized the cells to this drug, an effect that could be abolished by uridine co-treatment (Figure 5g). This phenotype was reproduced using VE-821 (Extended Data Figure 8g), and we also observed pyrimidine-dependent sensitization to BAY upon AA5 treatment in other transformed and non-transformed cell lines (Extended Data Figure 8h, i). Notably, ATR inhibitor sensitivity is not a general feature of aspartate-limited or slow-growing cells, as rotenone-treated cells in pyruvate-free media were not sensitized to BAY (Extended Data Figure 8j). Instead, our data indicate that this synergy was dependent on pyrimidine synthesis impairments, since aspartate, rotenone, or uridine treatment were all sufficient to abolish the toxicity of BAY and VE-821 in AA5 treated cells (Figure 5h, Extended Data Figure 8g). Collectively, these results suggest a model whereby SDH inhibition-induced succinate accumulation impairs pyrimidine biosynthesis, causing replication stress, S phase arrest, and sensitivity to interventions that prevent activation of replication stress signaling.

### SDH loss impairs ATCase activity in cells and tumors

To rule out any potential off-target effects of AA5 and test the effects of chronic SDH loss on pyrimidine synthesis, we used CRISPR/Cas9 to generate clonal SDHB-KO 143B cells^7^ (Figure 6a) that show depletion of fumarate and accumulation of succinate compared to parental cells, in accordance with impaired SDH activity (Figure 6b, Extended Data Figure 9a). In this system of chronic SDH impairment, there is no notion of an aspartate rebound. Instead, we would expect SDH-KO cells to exist in a ‘post-rebound’ state of low ATCase activity, impaired pyrimidine synthesis, and persistent replication stress. Indeed, SDH-KO cells have a lower carbamoyl-asp/aspartate ratio and decreased UMP levels compared to parental controls, consistent with ATCase impairment and resulting pyrimidine deficiency (Figure 6c, e, Extended Data Figure 9b-c).

**Figure 6.**
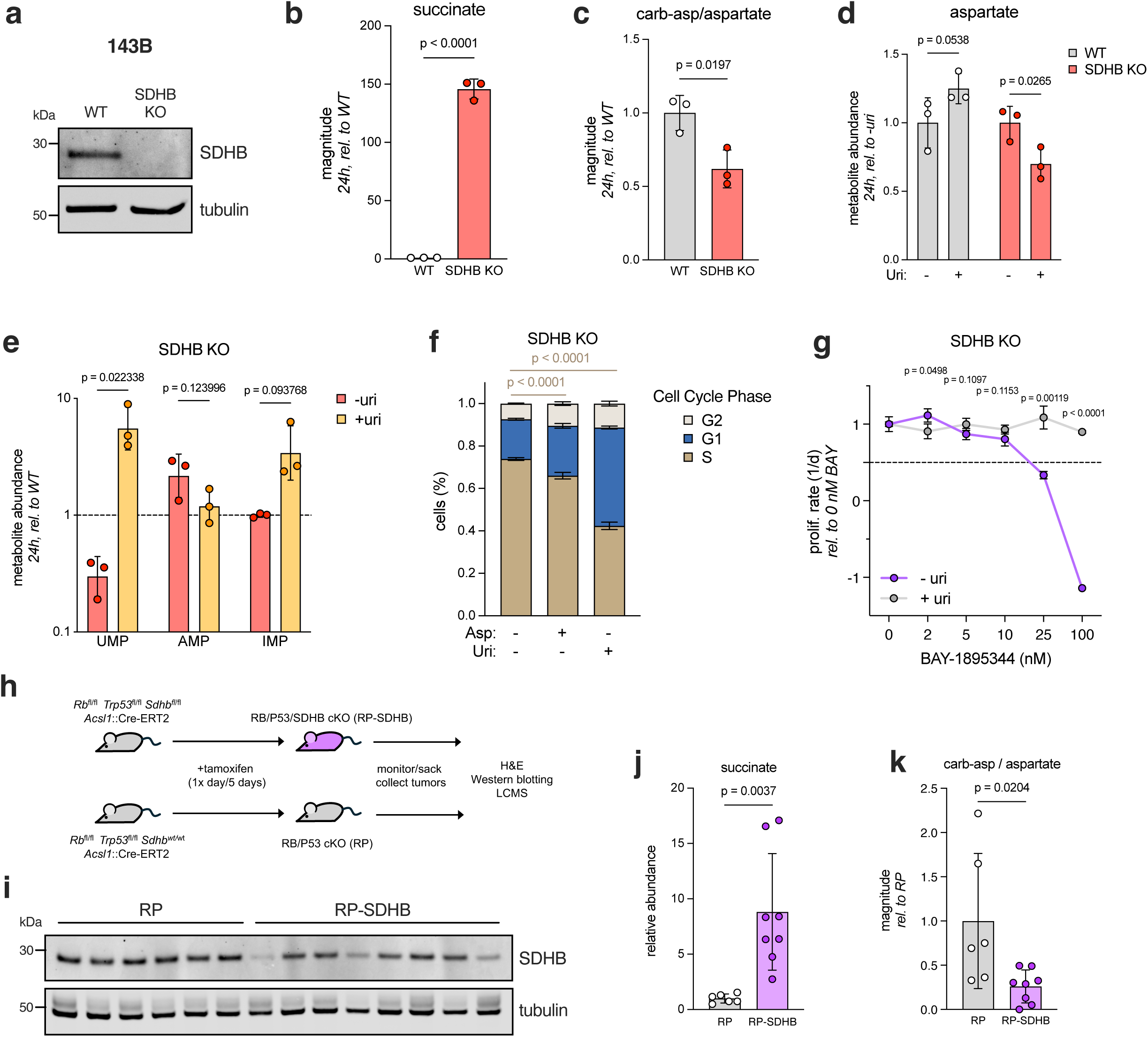
SDH loss impairs ATCase activity in cells and tumors. **a**, representative Western blot demonstrating levels of SDHB and tubulin loading control in SDHB-KO 143B cells. **b**, relative succinate abundances measured by LC-MS on SDHB-KO 143B cells 24 hours after media change (n=3). **c**, relative carbamoyl-aspartate/aspartate ratio measured by LC-MS in SDHB-KO 143B cells 24 hours after media change (n=3). **d**, relative aspartate abundances (normalized to untreated condition for each cell line) measured by LC-MS on wild-tye (WT) and SDHB-KO 143B cells 24 hours after treatment with vehicle control or 200 μM uridine (n=3). **e**, relative abundances of UMP, AMP, and IMP measured by LC-MS on SDHB-KO 143B cells 24 hours after treatment with vehicle control or 200 μM uridine (n=3). Abundances are normalized to those of wild type 143B cells in the same experiment, which is represented by the dotted line at y=1. **f**, representative cell cycle experiment showing the proportion of SDHB-KO 143B cells in each cell cycle phase 24 hours after treatment with vehicle control, 20 mM aspartate, or 200 μM uridine (n=3). **g**, proliferation rates (normalized to 0 nM BAY in each respective condition and measured using a conventional, 72-hour endpoint proliferation assay) of SDHB-KO 143B cells treated with vehicle control or 200 μM uridine and the indicated doses of BAY (n=3). **h**, schematic illustrating the design and experimental layout of the *Rb*^−/−^, *Tp53*^−/−^, *Sdhb*^−/−^ (RP-SDHB) mouse model. **i**, Western blot demonstrating levels of SDHB and tubulin loading control in individual *Rb*^−/−^, *Tp53*^−/−^, *Sdhb*^−/−^ (RP-SDHB) and littermate control (*Rb*^−/−^, *Tp53*^−/−^, RP) pituitary tumor extracts (n=6 RP, 8 RP-SDHB). **j**, relative succinate levels measured using LC-MS on RP and RP-SDHB tumor extracts in (i) (n=6 RP, 8 RP-SDHB). **k**, relative carbamoyl-aspartate/aspartate ratio measured using LC-MS on RP and RP-SDHB tumor extracts in (i) (n=6 RP, 8 RP-SDHB). Unless otherwise noted, experiments were conducted in DMEM with 1 mM pyruvate, data represented as mean +/− S.D. Statistical significance determined using unpaired t-tests (panels b-e, j-k) or an ordinary two-way ANOVA with uncorrected Fisher’s LSD and a single pooled variance (panels f-g). In panel f, asterisks denote results of statistical testing comparing the S-phase fraction in the indicated treatment conditions.

Given our previous results, we would also predict that supplementing uridine to pyrimidine-limited SDH-KO cells would reengage aspartate consumption, thereby depleting aspartate levels until they became limiting for purine synthesis (Extended Data Figure 5). Consistent with this model, uridine supplementation significantly lowers aspartate levels in SDH-KO cells but not parental controls (Figure 6d), and uridine-treated SDH-KO cells show evidence of emergent AMP deficiency and IMP accumulation (Figure 6e). Next, we analyzed cell size, cell cycling, and ATRi sensitivity to determine whether permanent SDH loss results in persistent replication stress. Strikingly, SDH-KO cells show significantly larger cell volumes than parental controls, an increased proportion of cells in S-phase, and sensitivity to BAY (Figure 6f, g, Extended Data Figure 9d). All three of these phenotypes could be partially or fully rescued by aspartate or uridine, supporting the hypothesis that genetic SDH loss causes replication stress via the mechanisms outlined above.

Finally, we sought to test the relationship between SDH function and ATCase activity in a more physiological setting. SDH subunits have been implicated as tumor suppressors for several neuroendocrine tumor types, including pituitary adenomas^53–60^. To this end, we generated mice in which the tumor suppressors *Rb*, *Trp53*, and *Sdhb* are simultaneously knocked out using tamoxifen-inducible Cre-ERT2 under the neuroendocrine-specific *Ascl1* promoter (Figure 6h). These mice reproducibly form pituitary adenomas over several months following tamoxifen induction, although SDHB loss did not produce a more aggressive disease in this system (Extended Data Figure 9f-g).

We harvested tumors at the end point and analyzed samples by both Western blotting and LC-MS to examine SDH loss and its effects on metabolic state (Figure 6h). Compared to pituitary tumors from *Rb*– and *Trp53*-null (RP) littermate controls, tumors from *Rb*-, *Trp53*-, and *Sdhb*-null (RP-SDHB) mice showed significantly lower, but not uniformly absent, SDHB protein levels, likely reflecting some mosaicism in the resulting tumors (Figure 6i, Extended Data Figure 9e). Nonetheless, RP-SDHB tumors collectively showed higher succinate levels than RP controls (Figure 6j), consistent with impaired SDH activity, which anticorrelated with SDHB protein abundance on a per-sample basis (Figure 6j, Extended Data Figure 9h). Next, we examined the carbamoyl-aspartate/aspartate ratio as a metabolic indicator of ATCase activity. Consistent with SDH loss inhibiting ATCase activity, the carbamoyl-aspartate/aspartate ratio was significantly lower in RP-SDHB tumors compared to RP controls (Figure 6k, Extended Data Figure 9i, j). On a per-sample basis, carb-asp/aspartate and succinate showed an exponential relationship, consistent with a threshold of succinate abundance above which ATCase activity becomes impaired (Extended Data Figure 9k).

Overall, our results suggest a cohesive model describing the interaction between SDH activity, pyrimidine synthesis, and aspartate abundance under both and acute and chronic settings of SDH impairment (Figure 7). Under basal cellular conditions, both SDH and ATCase are active, aspartate levels are high and succinate pools are low. Acute SDH impairment causes an immediate decrease in aspartate acquisition (production), while proliferation impairments lag metabolic changes, leading to a net decrease in aspartate levels. At some point, aspartate levels decrease and succinate accumulates enough to inhibit pyrimidine synthesis at ATCase, leading to pyrimidine deficiency, replication stress, and S-phase accumulation. In this regime, overall aspartate consumption dips even lower than its acquisition, leading to an aspartate rebound. As aspartate levels increase, however, they once again become competitive with succinate, licensing some ATCase activity/pyrimidine synthesis and temporarily releasing the cells from S phase. Cells now exist in a regime in which aspartate levels are kept ‘artificially’ high due to succinate-mediated ATCase inhibition: aspartate levels rise until ATCase is sufficiently disinhibited to support proliferation, after which aspartate is consumed and once again falls below the threshold needed to sustain ATCase activity, pyrimidine synthesis, and proliferation. Cells exhibiting chronic SDH impairment exist permanently in this last regime, characterized by persistent replication stress and slow cell cycling (Figure 7).

**Figure 7.**
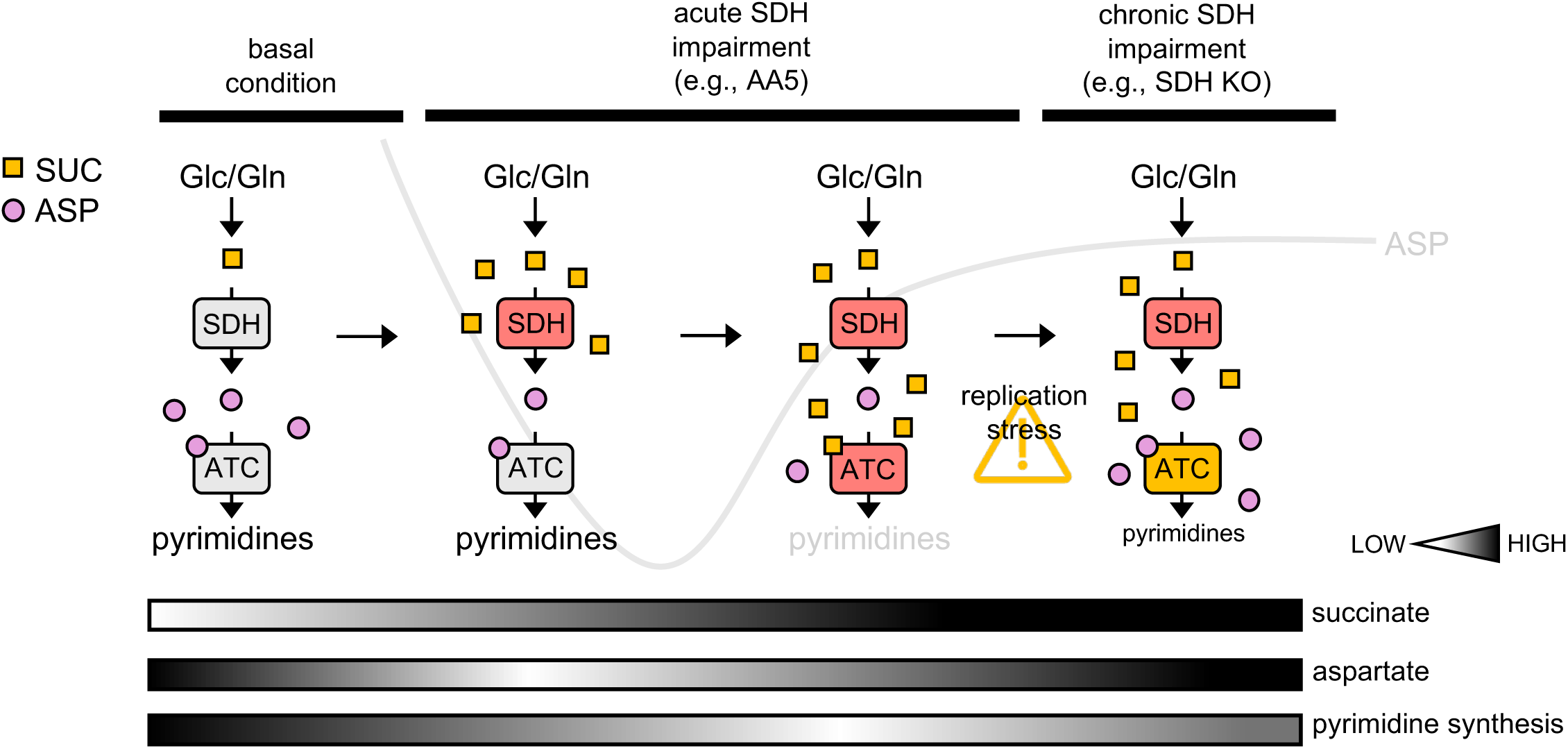
Model depicting the interaction between SDH status, metabolite levels, and ATCase activity.

## Discussion

Here, we leverage an aspartate biosensor and live cell imaging to exemplify a model in which cellular aspartate abundance dynamics are determined by its acquisition and consumption. The variable aspartate behaviors found under different metabolic contexts observed here also reinforce the principle that metabolite levels neither directly equate to pathway flux nor inherently identify how metabolic perturbations impact cell function, two common misconceptions in metabolism research^61^. In the case of aspartate limitation, we reveal that aspartate depletion causes functional effects once aspartate becomes limiting for the synthesis of specific anabolic fates. For SDH inhibition, we find a hierarchy of deficiencies for synthesis of pyrimidines, then purines, then proteogenic fates. Overall, these results highlight the power of nondestructive, temporally-resolved measurements to capture complex metabolic dynamics that are missed by traditional metabolomics measurements conducted at limited timepoints.

Our mechanistic investigation of the metabolic consequences following SDH deficiency reveals a novel role for succinate as a modulator of mammalian pyrimidine biosynthesis at ATCase. Since succinate accumulation is observed in physiological contexts beyond SDH-mutant cancers, including hypoxia, ischemia/reperfusion, liver inflammation, macrophage function, and brown adipose tissue thermogenesis^62–73,29^, it will be important to consider the potential roles that pyrimidine synthesis impairments play in these contexts, which may also depend on the availability of salvageable nucleotide precursors in the microenvironment^74–76^. Finally, our findings in SDH-KO cells and tumors raise the question of whether SDH-mutant tumors may be sensitive to interventions targeting the consequences of pyrimidine deficiency, including replication stress. Consistent with this possibility, several studies of preclinical models of SDH-deficient cancers have found evidence of alterations to nucleotide homeostasis and genomic integrity^30,77–79^. Overall, this work expands the growing understanding of how metabolites can regulate cell function beyond their direct roles as catabolic or anabolic substrates.

## Acknowledgments

We thank members of the Sullivan lab for discussion and feedback, Z. Rasmussen and J. Mayers of the Mayers lab at Fred Hutch for guidance and materials related to MBP-ATCase cloning and expression, J. Young and Y. Arimura of the Arimura lab at Fred Hutch for guidance and materials on MBP-ATCase purification, A. Kaiser and L. Doyle of the Stoddard lab at Fred Hutch for guidance and materials related to mass photometry and MBP-ATCase expression, B. Stoddard for discussion and feedback, and O.C. Young for moral support. This research was supported by the Proteomics & Metabolomics Shared Resource of the Fred Hutch/University of Washington Cancer Consortium (P30CA015704) and Fred Hutch Scientific Computing is supported by NIH grants S10-OD-020069, and S10-OD-028685. L.B.S. and D.M. acknowledge support from a pilot award from the Human Biology Division at Fred Hutch. L.B.S. acknowledges support from the NIGMS (R35GM147118) and the NCI (U54CA132381). D.S. was supported by PHS NRSA T32GM007270 from NIGMS.

## Contributions

M.L.H. and L.B.S. conceived of the project. M.L.H. and D.S. performed the experiments with assistance from S.D., E.Z., K.D., A.D.D., and D.M. L.B.S. supervised the project. M.L.H., D.S., and L.B.S. wrote the manuscript with input from all authors.

## Data Availability

All data supporting the findings of this study are available within the paper and its supplementary information.

For Extended Data Figure 4, relevant data, code, and analysis can be found at https://github.com/krdav/lab-work/tree/main/nucleotide_salvage_tracing.

## Supplementary Videos

Supplementary Video 1: Green and red channels overlaid for 143B sensor cells cultured in DMEM without Pyruvate.

Supplementary Video 2: Green and red channels overlaid for 143B sensor cells cultured in DMEM with 1 mM Pyruvate.

Supplementary Video 3: Green and red channels overlaid for 143B sensor cells cultured in DMEM without Pyruvate, following treatment with 50 nM rotenone.

Supplementary Video 4: Green and red channels overlaid for GOT1/2 DKO 143B sensor cells cultured in DMEM without Pyruvate, following a media change from 20 mM to 6 mM aspartate.

Supplementary Video 5: Green and red channels overlaid for 143B sensor cells cultured in DMEM with 1 mM Pyruvate, treatment with 5 µM AA5.

## Materials and Methods

### Cell Culture

Cell lines (143B, H1299, HCT116, 293T pLentiX, HT1080) were acquired from ATCC and regularly tested to be free from mycoplasma contamination (MycoProbe, R&D Systems). Cells were maintained in Dulbecco’s Modified Eagle’s Medium (DMEM) (Gibco, 50-003-PB) supplemented with 3.7 g/L sodium bicarbonate (Sigma-Aldrich, S6297), 10% fetal bovine serum (FBS) (Gibco, 26140079) and 1% penicillin-streptomycin solution (Sigma-Aldrich, P4333). Cells were cultured in a humidified incubator at 37°C with 5% CO_2_.

### Generation of jAspSnFR3/NucRFP cell lines

Nuclear RFP cell lines were generated as previously described^21^. jAspSnFR3 lentivirus was generated by co-transfection of HEK293T pLentiX cells with p-Lenti-jAspSnFR3 (Addgene, 203488) and the packaging plasmids pMDLg/pRRE (Addgene, 12251), pRSV-Rev, (Addgene, 12253) and pMD2.G (Addgene, 12259) using FuGENE transfection reagent (Fisher, PRE2693) in DMEM (Fisher, MT10017CV) without FBS or penicillin-streptomycin. The supernatant containing lentiviral particles was filtered through a 0.45 µM membrane (Fisher, 9720514) and was supplemented with 8 µg/µL polybrene (Sigma, TR-1003-G) prior to infection. For infection, 143B and GOT1/2 DKO 143B cells were seeded at 50% confluency in 6 well dishes and centrifuged with lentivirus (900g, 90 mins, 30°C). After 24 hours the media was replaced and after 48 hours cells were treated with 150 µg/mL hygromycin (Sigma-Aldrich, H7772-1G) and maintained in selection media until all uninfected control cells died. After selection, cells were expanded and single cell cloned by limiting dilution, plating 0.5 cells/well using two 96 well plates. These clones were incubated until 10-30% confluency and screened for high GFP and RFP signal using an Incucyte S3 (Sartorius). The highest expressing monoclonal cells were selected and further expanded on 6 well plates and again screened for high fluorescence using the Incucyte. From this, single clones were chosen, expanded and used for all subsequent experiments. H1299 and GOT1/2 DKO H1299 cells expressing jAspSnFR3 and nuclear RFP were previously generated^21^, whereas WT 143B, GOT DKO 143B, and cells were engineered to express nuclear RFP (Cellomics, PLV-10205-50) and pLenti-jAspSnFR3 for this manuscript. GOT1/2 DKO 143B and H1299 cells (no aspartate sensor) with and without SLC1A3 expression were also previously generated^21^.

### jAspSnFR3 and NucRFP Incucyte measurements

Experiments were conducted in either DMEM without pyruvate (Corning 50-013-PB) or DMEM with 1 mM pyruvate (Sigma, P8574) as indicated in the figure legends, supplemented with 3.7 g/L sodium bicarbonate, 10% dialyzed FBS (Sigma-Aldrich, F0392) and 1% penicillin– streptomycin solution. To start an experiment, cells were trypsinized (ThermoFisher, 25200056), resuspended in media, counted using a Coulter counter (Beckman Coulter, Multisizer 4) and seeded onto 24-well dishes (Nunc, 142475) with an initial seeding density of 15,000, 18,000, 50,000, or 20,000 cells/well for H1299, 143B, H1299/143B GOT1/2 DKO, respectively. After 24 hours of incubation, the plates were moved into an Incucyte S3 (Sartorius) live cell imaging platform inside a humidified incubator at 37°C with 5% CO_2_ for a pre-treatment scan. Once the scan was complete, plates were removed for treatment. Drug treatments such as Atpenin A5 (MedChemExpress, HY-126653) and Rotenone (Sigma-Aldrich, R8875) were spiked-in as 2x solutions in DMEM without pyruvate and dialyzed FBS along with 2x sodium pyruvate, where applicable. For treatments with varying media aspartate (Sigma-Aldrich, A7219) wells were washed with PBS and filled with DMEM containing the various aspartate concentrations. For plates receiving asparagine (Sigma-Aldrich, A7094) or uridine (Sigma-Aldrich, U3003), stocks were generated in water and made into 2x stocks in media, before a final dilution into experimental media so that the final concentration was 500 µM (Asn) or 200 µM (Uri), with vehicle wells receiving an equivalent volume of media with water in the place of the metabolite additives. Adenine (Sigma, A2786) was prepared fresh for each experiment by dissolving powder in 500 µL 1M HCl, neutralizing with 500 µL 1M NaOH, and filtering through a 0.22 µm filter (Fisher, FB12566506). This solution was then added to fresh media so that the final concentration was 100 µM. Live cell imaging was performed on the Incucyte S3 using the GFP and RFP channels with default exposure times. Images were processed using the associated Incucyte software to subtract background, define areas of cell confluence and GFP/RFP signal and extract integrated GFP/RFP values per well. Reported GFP/RFP values are normalized to the pretreatment scan on a per-well basis. Nuclei were counted as RFP instances at each timepoint, and average proliferation rates were calculated in 7-9 hour sliding windows according to the formula:

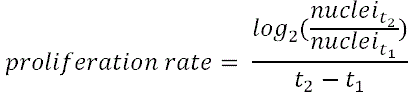

where t_1_ and t_2_ are initial and final scan times measured in days, respectively, and proliferation rate is calculated as cell doublings/day. Aspartate rebound was quantified by dividing the final normalized GFP/RFP value with the minimum normalized GFP/RFP value during an assay, per well.

### Conventional Proliferation Assays and Cell Volume Measurements

Cells were trypsinized, resuspended in media, and counted (Beckman Coulter Counter Multisizer 4 or Nexcelom Auto T4 Cell-o-meter) and seeded overnight onto 24-well dishes (Corning, 3516) with similar initial seeding densities described above. After overnight incubation, 3 wells were counted for a starting cell count at the time of treatment. Cells were washed in phosphate-buffered saline (PBS) and 1-2 mL of treatment media was added to each well. Experiments were conducted in DMEM without pyruvate supplemented with 3.7 g/L sodium bicarbonate 10% dialyzed FBS and 1% penicillin-streptomycin solution, with or without 1 mM sodium pyruvate, 20 mM aspartate, or 0.5 mM Asparagine, 200 µM uridine, 100 µM adenine, or 10-25 mM succinic acid (Sigma, S3674). Drug treatments included rotenone (Sigma-Aldrich, R8875), cycloheximide (Sigma, C7698), Atpenin A5 (MedChemExpress, HY-126653), BAY-1895344 (Selleckchem, S8666) and DMSO as a vehicle (Sigma, D2650). Cells were incubated in a humidified incubator at 37°C with 5% CO_2_ then counted after 2-4 days. At the endpoint, cells were counted using a Beckman Coulter counter Multisizer 4 instrument, which measures individual particle (cell) volumes in addition to counting cells. Proliferation rate was determined as described previously^7^.

### Generation of KO cells

Knockout cell lines were generated as previously described^7,21^. Three single guide RNA (sgRNA) sequences targeting fumarate hydratase (FH) were purchased (Synthego) and are listed in the table below. Each sgRNA was resuspended in nuclease-free water, combined with SF buffer (Lonza, V4XC-2032), and sNLS-spCas9 (Aldevron, 9212). 2×10^5^ 143B cells were resuspended in the resulting solution containing ribonucleoprotein complexes (RNPs) and electroporated using a 4D-Nucleofector (Amaxa, Lonza) using electroporation program FP-133. Nucleofected cells were then moved to a 12-well plate (Corning, 3513) and, after achieving confluence, were single-cell cloned by limiting dilution by plating 0.5 cells/well in a 96 well plate. Gene knockout was confirmed using western blots on the nucleofected pool and each single cell clone used in this study. GOT1/2 DKO cells were generated in the same way and were previously described^21^.

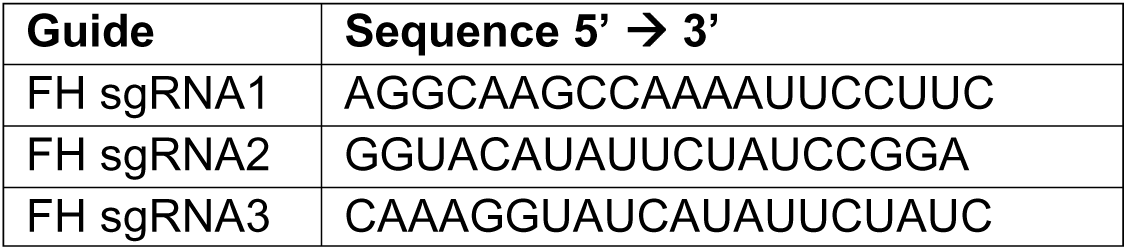

### Western Blotting

Protein lysates were harvested in RIPA buffer (Sigma, R0278) supplemented with protease inhibitors (Fisher, A32953) and phosphatase inhibitors (Fisher, 78442). Protein concentration was determined using a Bicinchoninic Acid Assay (Fisher, 23225) using bovine serum albumin (BSA) as a protein standard. Equal amounts of protein were denatured with Bolt 4x Loading Dye (ThermoFisher, B0007) and Bolt 10x reducing agent (ThermoFisher, B0004), heated at 95°C for 2-5 min, and loaded onto 4-12% by SDS-polyacrylamide gels (Invitrogen, NW04127). After electrophoretic separation, proteins were transferred onto a 0.22 mm nitrocellulose using iBlot2 transfer stacks (Fisher, IB23001) and transferred with the P0 system setting. Membranes were blocked with 5% BSA in Tris-buffered saline with 0.1% Tween-20 (TBS-T) and incubated at 4°C overnight with the following antibodies: anti-GOT2 (Proteintech, 14800-1-AP, 1:1000), anti-GOT1 (Cell Signaling, 34423S, 1:1,000), anti-GFP (Sigma, 1:1000), anti-FH (Origene, TA500675S, 1:1000), anti-SDHB (Atlas, HPA002868, 1:1,000), anti-pChk1 (Cell Signaling, 2348S, 1:1000), anti-Chk1 (Cell Signaling, 2G1D5, 1:1000), anti-pChk2 (Cell Signaling, 2197S, 1:1000), anti-GAPDH (Cell Signaling, 5174S, 1:5000), and anti-Vinculin (Sigma, SAB4200729, 1:10,000), anti-tubulin (Sigma, T6199). The next morning, membranes were washed three times with TBS-T, and the following secondary antibodies were added: 800CW Goat anti-Mouse IgG (LiCOR, 926-32210; 1:15,000), 680RD Goat anti-Rabbit IgG (LiCOR, 926-68071; 1:15,000). Membranes were washed three more times with TBS-T and imaged on a LiCOR Odyssey Near-Infrared imaging system.

### RNA Extractions and tRNA Aminoacylation Quantification

143B cells were grown on 6 well plates in DMEM without pyruvate, in dialyzed FBS. For the Atpenin A5 treatment, 1 mM pyruvate was added to the media. At 50% confluency, cells were treated with drugs in replicates for 30 hours. 143B cells were treated with vehicle (DMSO), rotenone (50 nM), or Atpenin A5 (5 μM). For RNA extraction, the media on the cells was quickly and thoroughly aspirated before adding 3 mL TRIzol (ThermoFisher, A33250) to cover all the cells. From this point onward, everything was kept ice-cold to prevent hydrolysis of the aminoacylation. After a 2 min incubation, the cell material was scraped down the slope mixing it with the TRIzol, then 2×1.5 mL was transferred to 2 mL Eppendorf tubes and 0.3 mL chloroform was added. The tubes were vortexed 2 min and then centrifuged (17,000×*g*, 5 min). From each tube, 0.75 mL of the upper layer was transferred to a tube with 0.8 mL isopropanol (IPA) (ThermoFisher, A464SK), then mixed and incubated 60 min at –20°C. Tubes were then centrifuged (17,000×*g*, 15 min) and RNA pellets were washed twice with 1 mL 80% IPA containing 100 mM sodium acetate (pH = 4.5) (Sigma, S7545). Washes are critical to prevent glycerol present in TRIzol from inhibiting the subsequent periodate oxidation step. A last wash was performed using 1 mL 100% IPA and after removing the supernatant the RNA pellets were air-dried at room temperature, then stored dry at –80°C. Aminoacylation levels, referred to as “charge”, were measured by sequencing and determining the fraction of tRNAs protected from periodate oxidation and 3’ nucleotide cleavage, as previously described in detail^24^. The values for each of the multiple tRNA transcripts decoding each codon were calculated as an expression weighted average of all codon-specific transcript charges.

### Polar Metabolite Extractions

<u>Time-course</u>. For LC-MS measurements across several time-points, 143B cells were seeded overnight at either 2×10^5^, 1×10^5^, or 0.5×10^5^ cells per well of a 6-well dish for the 10/16hr, 24hr, and 44hr time points, respectively. The next day, cells were washed twice with PBS and changed to the indicated medias supplemented with 1 mM pyruvate, 10% dialyzed FBS, 1% penicillin-streptomycin, and treatments as indicated, and returned to the tissue culture incubator. Polar metabolites were extracted from cells by three washes with ice-cold blood bank saline, (Fisher, 23293184) then 300 µL of 80% HPLC grade methanol in HPLC grade water was added to each well and cells were scraped with the back of a P1000 pipet tip and transferred to Eppendorf tubes. Tubes were centrifuged (17,000xg, 15 mins, 4°C) and the supernatant containing polar metabolites was transferred to a new centrifuge tube and placed in a Centrivap until lyophilized. Corresponding plates with the same treatment conditions were trypsinized at the same time metabolites were extracted and counted on a Beckman Coulter Counter to obtain total cell volume per well. Average cell volumes for each condition were calculated and used to normalize metabolites after centrifugation. Dried metabolites were resuspended in 40 µL solvent per µL cell volume in 80% HPLC grade MeOH containing ^13^C-labeled metabolites made by partial hydrolysis (12 hours in 6 M HCl at 90°C) of U-^13^C spirulina whole cells lyophilized powder (Cambridge Isotope Laboratories, CLM-8400-PK) as an internal standard to account for matrix effects and absolute quantitation of intracellular metabolites. Ion counts were normalized to the internal standard metabolite when appropriate to determine a response ratio. *<u>Succinate and Aspartate Concentration Measurements.</u>* A standard curve of known succinate and aspartate concentrations over three orders of magnitude was generated in 80% HPLC-grade MeOH containing ^13^C spirulina standard and run in parallel with the time-course experiment detailed above. Response ratios for succinate and aspartate ion counts from cell extracts were used to calculate a concentration according to the standard curve, which were then back calculated by cell volume to generate intracellular concentrations. *<u>Other Measurements.</u>* For standard metabolic analysis, cells were seeded overnight at 1×10^5^ cells per well of a 6-well dish. The next day, cells were washed twice with PBS and changed to the indicated medias supplemented with 10% dialyzed FBS, 1% penicillin-streptomycin, and treatments as indicated, and returned to the tissue culture incubator. After 30-32 hours, polar metabolites were extracted from cells by the same protocol as mentioned above.

### Isotope Tracing

*^15^N Glutamine tracing.* The fractional contribution of individual components into their respective aspartate consuming fate was determined in 143B and H1299 cells for the salvageable metabolites asparagine (Asn), uridine (Uri), adenine (Ade), hypoxanthine (Hpx) (Cayman, 22254), and guanine (Gua) (Sigma, G11950), along with a vehicle treatment (Vec). The salvageable metabolites were spiked in from a 20x stock solution to achieve a final concentration of: 500 μM Asn, 200 μM Uri, 100 μM Hpx, 100 μM Ade or 100 μM Gua. The fraction of salvage was determined by stable isotope tracing, performed using both Gln amide-^15^N (Cambridge Isotope Laboratories, NLM-557-PK) and Gln alpha-^15^N (Cambridge Isotope Laboratories, NLM-1016-PK) in separate reactions and added to DMEM without glucose, glutamine, pyruvate and phenol red (Sigma, D5030) supplemented with 10% dialyzed FBS, 1x penicillin-streptomycin and 25 mM glucose (Sigma, G7528). This combination of cell lines, salvageable metabolites, and tracers gave 24 conditions which were labelled to steady state by culturing for four passages with a 1/20 split at each passage. At the end of the last passage each condition was split into four technical replicates and plated on 24 well plates. Upon reaching confluency, polar metabolites were extracted and submitted to LC-MS. The relative contribution of guanine salvage into the GTP pool was determined using the m+0 vs. m+3 GTP isotopologue fractions from the Gln amide-^15^N labelled samples. The relative contribution of both adenine and hypoxanthine salvage into the ATP pool was determined using the m+0 ATP isotopologue fraction from the Gln amide-^15^N labelled samples and the direct contribution from adenine was determined using the m+1 isotopologue fractions of aspartate compared to ATP from the Gln alpha-^15^N labelled samples. *U-^13^C Glutamine tracing.* WT 143B cells were seeded at 1×10^5^ cells per well of a 6-well dish. The next morning, cells were washed twice with PBS and changed to DMEM without glucose, glutamine, pyruvate, or phenol red (Sigma, D5030) supplemented with 10% dialyzed FBS, 1% penicillin-streptomycin, 1 mM pyruvate, 25 mM ^12^C glucose (Sigma, G7528), and 4 mM U-^13^C glutamine (Cambridge Isotopes, CLM-1822). 143B cells were treated as indicated for 32 hours, then harvested by the same protocol detailed above.

### Liquid Chromatography-Mass Spectrometry (LC-MS)

Resuspended samples were transferred to liquid chromatography-mass spectrometry (LCMS) vials for measurement by LCMS. Metabolite quantitation was performed using a Q Exactive HF-X Hybrid Quadrupole-Orbitrap Mass Spectrometer equipped with an Ion Max API source and H-ESI II probe, coupled to a Vanquish Flex Binary UHPLC system (Thermo Scientific). Mass calibrations were completed at a minimum of every 5 days in both the positive and negative polarity modes using LTQ Velos ESI Calibration Solution (Pierce). Polar Samples were chromatographically separated by injecting a sample volume of 1 μL into a SeQuant ZIC-pHILIC Polymeric column (2.1 x 150 mm 5 mM, EMD Millipore). The flow rate was set to 150 μL/min, autosampler temperature set to 10 °C, and column temperature set to 30 °C. Mobile Phase A consisted of 20 mM ammonium carbonate and 0.1 % (v/v) ammonium hydroxide, and Mobile Phase B consisted of 100 % acetonitrile. The sample was gradient eluted (%B) from the column as follows: 0-20 min.: linear gradient from 85 % to 20 % B; 20-24 min.: hold at 20 % B; 24-24.5 min.: linear gradient from 20 % to 85 % B; 24.5 min.-end: hold at 85 % B until equilibrated with ten column volumes. Mobile Phase was directed into the ion source with the following parameters: sheath gas = 45, auxiliary gas = 15, sweep gas = 2, spray voltage = 2.9 kV in the negative mode or 3.5 kV in the positive mode, capillary temperature = 300 °C, RF level = 40 %, auxiliary gas heater temperature = 325 °C. Mass detection was conducted with a resolution of 240,000 in full scan mode, with an AGC target of 3,000,000 and maximum injection time of 250 msec. Metabolites were detected over a mass range of 70-1050 *m/z*. Quantitation of all metabolites was performed using Tracefinder 4.1 (Thermo Scientific) referencing an in-house metabolite standards library using ≤ 5 ppm mass error. Data from stable isotope labeling experiments includes correction for natural isotope abundance using IcoCor software v.2.2.

For experiments in Extended Data Figure 7a-d, polar metabolite extracts were generated as described above and resuspended in HPLC-grade 80% methanol without stable isotope standards. Metabolite quantitation was performed using an Agilent 6495D Triple-Quadrupole Mass Spectrometer equipped with an Agilent JetStream Heated ESI source coupled to an Agilent 1290 Infinity II UHPLC (Agilent Technologies). A Checktune was performed using the Agilent ESI-L low concentration tuning mix to assess the status of the instrument prior to data collection. Polar Samples were chromatographically separated by loading a sample volume of 2μL onto one of two Poroshell 120 HILIC-Z columns (2.1×150mm 2.7micron, Agilent Technologies) running in a bespoke alternating column regeneration method. The flow rate was set to 600 μL/min, multisampler temperature set to 4 °C, and column temperature set to 25 °C. Mobile Phase A consisted of 0.1% formic acid with 10mM ammonium formate, and Mobile Phase B consisted of acetonitrile with formic acid at 0.1% (v/v). The sample was gradient eluted (%B) from the column as follows: 0-0.14min, initial hold at 95% “B”; 0.14-2.29 min, 95% to 40% “B”; 2.29-4.0 min, hold at 40% “B”. At the end of the method, the column oven valve would switch over to the other (K’ value matched) HILIC-Z column and the gradient would run on column 2, while column 1 was regenerated at 40% to 95% “B” from 0-0.56 min, followed by an increase in flow-rate to 1200 μL/min and held at 95% “B” until 10 column volumes were pumped through the column. Samples were analyzed on the mass spectrometer using the following parameters: gas flow = 13.0L/min, sheath gas = 11L/min, nebulizer = 35 psi, gas temperature = 200C, sheath gas temperature = 250 C, capillary voltage = 3000V, nozzle voltage = 1500V and a CAV voltage of 5V. Metabolites were targeted in the MRM mode with a dwell time of 20ms at unit resolution; compound collision energies (CEs) were previously optimized in the automated mode of MassHunter’s built in optimizer module (ver. 12.1). Quantitation of all metabolites was performed using MassHunter’s Quantitative Analysis module referencing an in-house metabolite standards library.

### Molecular Docking Studies

Molecular docking was performed on the Fred Hutch high performance computing cluster using Autodock Vina v1.2.7^80,81^ using the vina scoring function and exhaustiveness = 64. Receptors were prepared using ChimeraX 1.10.1 and the OpenBabel web server^81^ and docking was performed in a 15×15×15 Å grid box centered on a single CP/aspartate binding pocket with grid space = 0.375. To verify the accuracy of the docking approach, the ATCase inhibitor PALA was re-docked onto a crystal structure of PALA-bound human ATCase (PDB: 5G1N) and manually verified to agree with the experimentally determined pose (data not shown). Docking results were ranked and visualized using ChimeraX 1.10.1.

### ATCase Expression and Purification

The ATCase domain of human CAD was expressed and purified using a modification of a previously published workflow^82^. Briefly, a gBlock was designed and purchased containing residues 1915–2225 of the human CAD protein (cDNA from transcript ENST00000264705.9) separated from an N-terminal 6xHis-tagged maltose binding protein (MBP) by a TEV protease cleavage site, with an SSG linker separating the 6xHis tag from MBP, two SSG linkers immediately 5’ of the TEV site, and a GS linker immediately 3’ of the TEV site. Gibson Assembly primers were designed using NEBuilder (New England Biolabs) and used to amplify and incorporate Gibson overhangs onto the backbone and insert using Q5 hi-fidelity PCR mastermix (NEB, M0492).

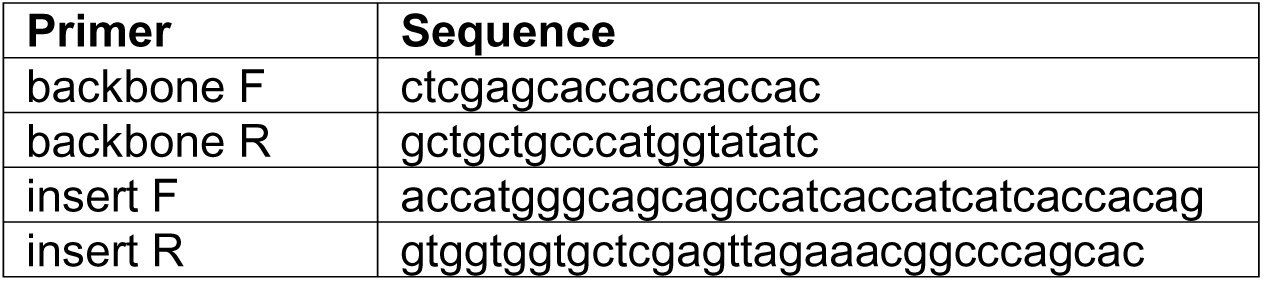

This construct was assembled into a pET-28a(+) backbone using GeneArt Hifi Gibson Assembly Mastermix (Invitrogen, A46627), transformed into TOP10 competent cells (Invitrogen, C404003), purified using miniprep/midiprep kits (Qiagen) and verified by long-read sequencing (Plasmidsaurus).

For protein expression, the MBP-ATCase construct was transformed into BL21 competent cells (Fisher, EC0114) and grown in autoinduction media^83^ at 37°C for 6 hours, then 18°C for 24 hours to induce protein expression. Bacterial pellets were sonicated at 70% amplitude, 2 sec on-5 sec off for 5 minutes on ice in lysis buffer (20 mM Tris-HCl pH 7.2, 500 mM NaCl, 10 mM imidazole, 5% glycerol, 0.5 mg/ml lysozyme (Thermo, 89833), 0.5% Octyl-B-D-thrioglucopyranoside (CHEM-IMPEX, 21810), 2 mM PMSF (Roche), 2 mM beta-mercaptoethanol (Sigma, M3148), adjust pH to 8.0 with NaOH), after which benzonase (∼25 U/ml,), MgCl_2_ (1 mM), and CaCl_2_ (1 mM) were added and lysate was incubated on ice for 15 minutes. Lysate was cleared by ultracentrifugation and batch-bound to HisPur Cobalt Resin (Thermo, 89964) equilibrated with binding buffer (20 mM Tris-HCl pH 7.2, 500 mM NaCl, 10 mM imidazole, 5% glycerol, adjust pH to 8.0) for 30 minutes in the cold room with rocking. After binding, resin was moved into a gravity column, washed 3×2 C.V. with wash buffer (20 mM Tris-HCl pH 7.2, 500 mM NaCl, 30 mM imidazole, 5% glycerol, 2 mM beta-mercaptoethanol, adjust pH to 8.0), and eluted with 5×1 C.V. elution buffer (wash buffer with 300 mM imidazole). MBP-ATCase presence and affinity purification was confirmed by SDS-PAGE of appropriate fractions on Bolt 4-12% Bis-Tris Plus polyacrylamide gels (Invitrogen) and subsequent Coomassie staining. Following affinity purification, elute fractions containing MBP-ATCase were pooled, concentrated to <2mL using a 10,000 kDa MW-cutoff spin column (Amicon, UFC901024), and loaded onto a Superdex 200 16/600 column (Cytiva) equilibrated in gel filtration buffer (20 mM Tris-HCl pH 7.2, 100 mM NaCl, 2% glycerol, 0.2 mM TCEP (Sigma, 646547)) for size exclusion chromatography (SEC) on an ÅKTA machine. Following SEC, another diagnostic protein gel was run, and the fractions indicated in Extended Data Figure 6c were pooled and stored at 4°C for use in downstream analyses. Final protein concentrations were estimated using spectrophotometry (Nanodrop).

While initial efforts attempted to cleave off the 6xHis-MBP tag using overnight dialysis in buffer containing TEV protease, it was found that isolated ATCase was unstable in solution. This fact—and the fact that ATCase has very few aromatic residues facilitating accurate detection and quantification by UV absorbance or BCA assays^82^—led to the decision to use the MBP-ATCase fusion protein in subsequent studies.

### ATCase Activity Assays

A colorimetric ATCase activity assays was established with minor modifications from previous studies^38,84^. Antipyrine/H_2_SO_4_ (5 g/L antipyrine (Sigma, A5882) in 50% v/v sulfuric acid (Sigma, 258105)) and oxime (8 g/L butanedione monoxime (Sigma, 31550) in 5% v/v glacial acetic acid (Millipore, K52336663013)) reagents were prepared and stored according to a previous study^84^. Briefly, purified MBP-ATCase was diluted in cold activity assay buffer (50 mM Tris-acetate pH 8.3, 0.01 mg/mL bovine serum albumin fraction V fatty acid free (Roche, 03117057001)) at ∼0.12 μΜ. Enzyme was portioned out into a 96-well plate at 50 μL/well, mixed with appropriate substrates (aspartate, succinate, and/or fumarate) diluted to 3X final concentration in activity assay buffer, and pre-incubated on the benchtop for 5 minutes. Reactions were started by rapidly adding 50 μL/well CP diluted to 3X final concentration in activity assay buffer, and allowed to proceed for 5 min in a benchtop incubator set to 37°C. After incubation, reactions were quenched by addition of 150 μL/well color solution (2:1 antipyrine/H_2_SO_4_: oxime reagents, prepared immediately before use), sealing the plate with plate film (Thermo), pulse vortexing, and heating on top of a 95°C heat block for ∼30min – 1 hour in room light. Plates were cooled to room temperature, and absorbance at 460 nm was subsequently quantified using a plate reader (Tecan). Blank controls were used to subtract background absorbance, and carbamoyl-aspartate calibration curves were run in each activity assay to verify linearity of the measured absorbances.

### Mass Photometry

Mass photometry was performed on a Two^MP^ instrument (Refeyn) similarly to what was previously described^85^ using AcquireMP v2024 and DiscoverMP v2024 software for acquisition and analysis, respectively. MBP-ATCase (∼40 nM) and appropriate substrates were prepared in mass photometry buffer (50 mM Tris-acetate pH 8.3) on ice and incubated for 2-4 minutes before imaging on uncoated glass slides (Refeyn). Droplet dilution autofocus was used to find focus in 20 μL droplet sizes. Mass calibrations were performed before each instrument run using MassFerence P1 calibrant (Refeyn, MP-CON-41033) according to manufacturer specifications. Multiple 1-minute movies were collected per sample and verified to agree qualitatively, and Gaussian fits were computed using DiscoverMP.

### Puromycin Incorporation Assay

143B cells were seeded in a 6-well plate at 200K cells/well. The following day, cells were washed two times with Dulbecco’s PBS (DPBS) and switched into appropriate treatment medias (2 mL/well, DMEM + 10% dialyzed FBS) for 24 hours. Puromycin incorporation was conducted by spiking in puromycin (Sigma, P9620) at 10 μg/mL for exactly 30 minutes before cells were washed once again and protein lysates were harvested in RIPA buffer (Sigma, R0278) supplemented with protease inhibitors (Fisher, A32953) and phosphatase inhibitors (Fisher, 78442). 100 μg/ml cycloheximide (Cayman, 14126) was added to the appropriate samples 30 minutes prior to puromycin addition. Protein concentrations were determined and Western blotting was performed as above. Membranes were incubated at 4°C overnight with the following antibodies: anti-puromycin (Kerafast, Kf-Ab02366-1.1, 1:1000) and anti-GAPDH (Cell Signaling, 5174S, 1:5000) and imaged as above. Densitometry on entire puromycin lanes was performed using ImageJ2 version 2.9.0.

### Cell Cycle Analysis

For cell cycle analysis, cells were plated in 6-well plates at 100K cells/well and incubated overnight. The following day, cells were washed three times with Dulbecco’s PBS (DPBS) and switched into appropriate treatment medias (3 mL/well, DMEM + 10% dialyzed FBS) for the indicated times. To fix cells, replicate wells were trypsinized, pelleted and washed twice with DPBS. Cells were resuspended in 300 μL ice-cold PBS, and 700 µL ice-cold 100% ethanol was added dropwise to each sample while vortexing to fix. Fixed cells were stored at –20°C until being processed for flow cytometry (no longer than 4 days). After all timepoints were collected, fixed cells were pelleted and washed with DPBS twice, then resuspended in 250 μL of 50 μg/mL propidium iodide (Biotium, 40017) with 100 μg/mL RNase A (Qiagen) staining solution for 1 hour at room temperature or overnight at 4°C, protected from light. Samples were then passed through a 0.35-μm filter into flow cytometry tubes (Falcon) before being analyzed on a BD FACSymphony A52 Cell Analyzer running FACSDiva software. 10,000 events were recorded per sample. Data was analyzed using the ‘Cell Cycle’ analysis module of FlowJo 10.10.1.

### Animal Studies

The *Rb1^lox/lox^; Trp53^lox/lox^; Ascl1-Cre-ERT2* model of neuroendocrine pituitary tumorigenesis, as described^86^ were bred to a *Sdhb^lox/lox^*allele from Dr. J. Favier to generate compound mutant mice used for this study^32^. Tamoxifen induction of Cre-ERT2 under the control of *Ascl1* promoter was accomplished by intraperitoneal injection of mice with tamoxifen (150 mg/kg per day), prepared in sterile corn oil, over five consecutive days. Mice were euthanized when they exhibited poor body condition and necropsies were performed, with the skull either fixed in Bouin’s fixative, or bisected, with part of the pituitary tumor excised and snap frozen on dry ice and the remainder fixed in the skull with Bouin’s for 1 week before processing and paraffin embedding. Tissue sections (4 μM thick) were stained with hematoxylin and eosin (H&E). All animal procedures were approved by the Institutional Animal Care and Use Committee (IACUC) at the Fred Hutchinson Cancer Center (protocol #50783).

### Mouse Pituitary Tumor Metabolomics and Western Blotting

For LC-MS analyses, frozen tumor chunks from RP/RP-SDHB mice were pulverized using liquid nitrogen-cooled instruments and portioned into pre-weighed tubes. HPLC-grade 80% MeOH was added at 1 mL/ 20 mg tissue and samples were vortexed for 30 mins at room temperature to extract polar metabolites. Samples were spun at 17,000 xg for 15 minutes in a refrigerated centrifuge, and 100 μL of supernatant was dried down in a refrigerated vacuum centrifuge overnight (Centrivap). Each sample was reconstituted in 500 μL of a 1:1 mix of ^13^C-labeled yeast and spirulina extracts (see above) and LC-MS was performed using a Q Exactive HF-X Hybrid Quadrupole-Orbitrap Mass Spectrometer as above. Wherever possible, ^13^C-labeled internal standards were used to calculate response ratios and correct for matrix effects. Due to systemic differences in global metabolite ion counts across samples, a ‘correction factor’ was created, consisting of the mean of the response ratios for five amino acids (leucine, lysine, threonine, tyrosine, and phenylalanine), and ion counts for metabolites investigated in Figure 6 were subsequently normalized using this factor. While this approach has been used in LC-MS studies previously^4^, we note that the carb-asp/aspartate ratio is insensitive to these systemic signal differences and is identically different between RP and RP-SDHB extracts whether or not this correction factor is used.

For Western blotting, cell material pellets leftover after the spin step above were briefly dried in a refrigerated vacuum centrifuge to remove residual extraction solvent, and 200 μL RIPA buffer with protease/phosphatase inhibitors and 5 mM EDTA was added per sample. Samples were vortexed for 5 min. at room temperature and incubated on ice for 15 min., then this vortexing/incubation procedure was repeated once more. Protein concentrations were determined and Western blotting was performed as above. Densitometry was performed using ImageJ2 version 2.9.0.

## Data Analysis

All graphs and statistical analyses were made in GraphPad Prism 10.4.1. Technical replicates, defined as parallel biological samples independently treated, collected, and analyzed during the same experiment, are shown. Experiments were verified with ≥ 2 independent repetitions showing qualitatively similar results. Details pertaining to all statistical tests can be found in the figure legends.

## Figure Legends

**Extended Data Figure 1.**
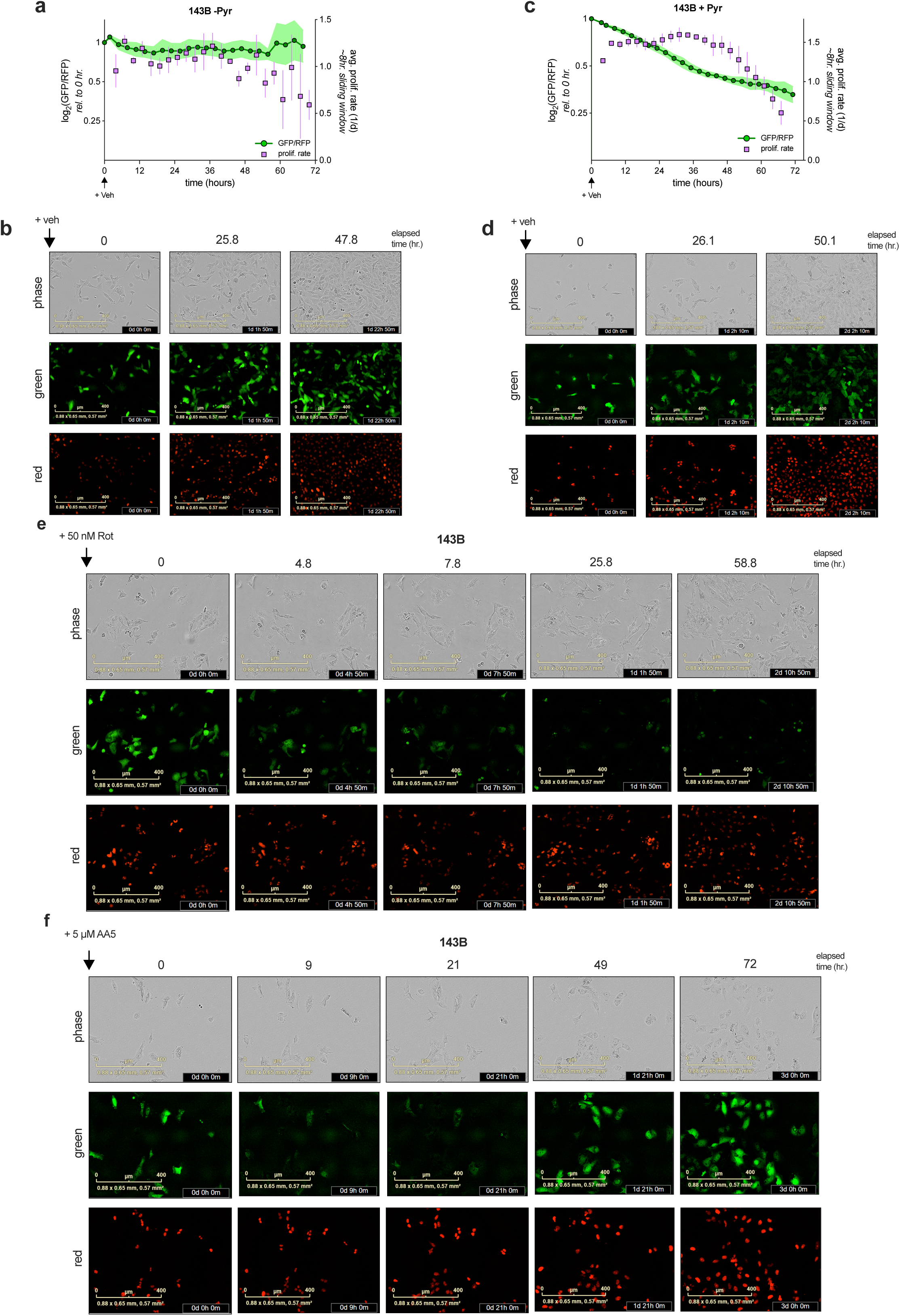
Related to figure 1. **a**, relative aspartate levels (green line, left y-axis) and absolute proliferation rates (purple points, right y-axis) of 143B cells in DMEM without pyruvate treated with DMSO (n=4). **b**, representative images from panel (a) taken at 0-, 25.8-, and 47.8-hours post-treatment. Phase contrast, green (jAspSnFR3), and red (nucRFP) channels are shown. **c**, relative aspartate levels and absolute proliferation rates of 143B cells in DMEM with 1 mM pyruvate treated with DMSO (n=3). **d**, representative images from panel (c) taken at 0-, 26.1-, and 50.1-hours post-treatment. Phase contrast, green, and red channels are shown. **e**, representative images from the live-cell imaging experiment in Figure 1b taken at five timepoints post-treatment. Phase contrast, green, and red channels are shown. **f**, representative images from the live-cell imaging experiment in Figure 1d taken at five timepoints post-treatment. Phase contrast, green, and red channels are shown. Data represented as mean +/− S.D.

**Extended Data Figure 2.**
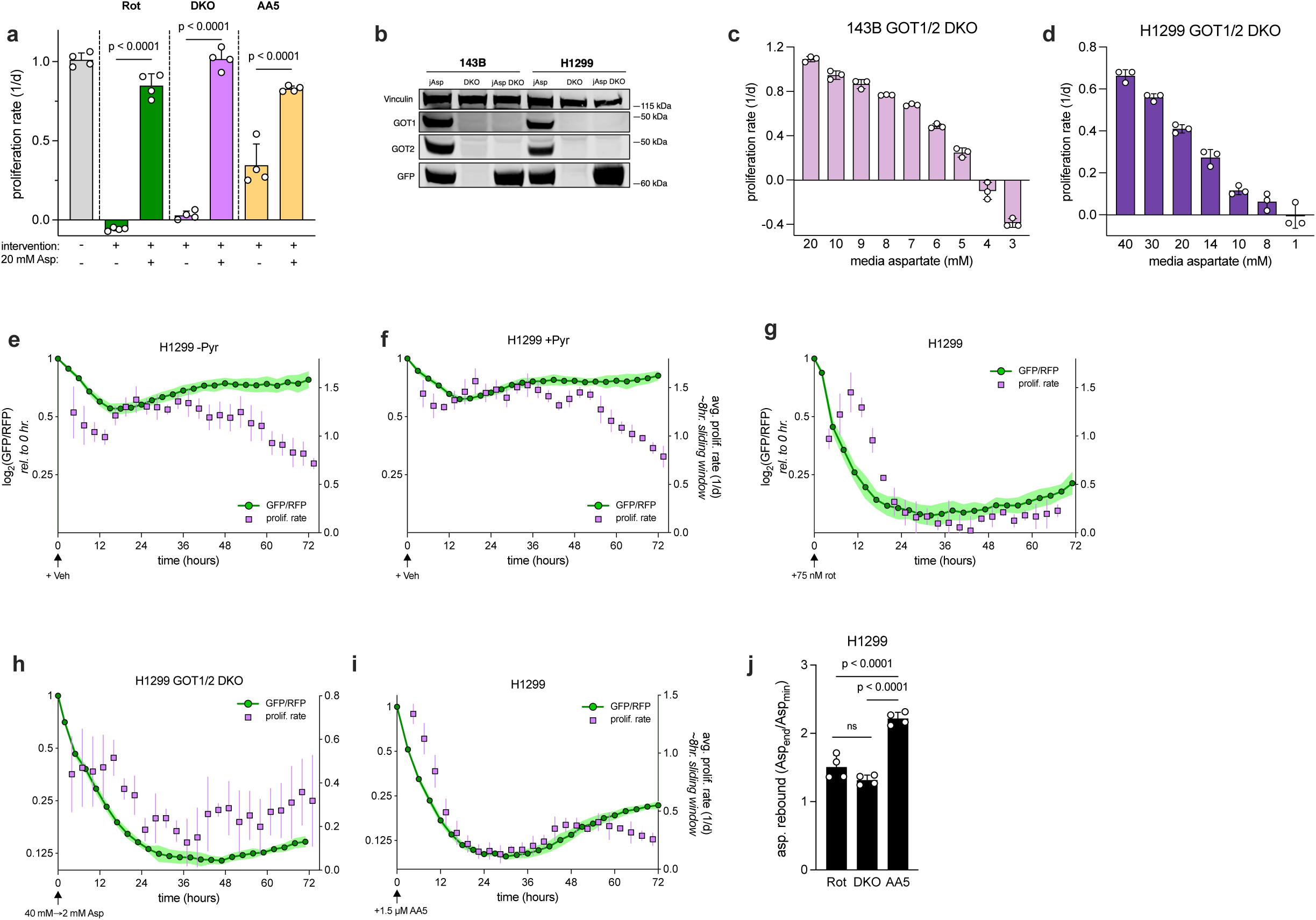
GOT1/2 DKO validation and aspartate/proliferation measurements in H1299 cells. **a**, average proliferation rates (determined using a conventional, endpoint proliferation assay) of 143B jAspSnFR3/NucRFP or GOT1/2 DKO 143B jAspSnFR3/NucRFP cells under the three paradigms of aspartate limitation [50 nM rotenone (Rot), aspartate withdrawal (5 mM) in GOT1/2 DKOs (DKO), and 5 µM Atpenin A5 (AA5)], with or without 20 mM aspartate (n=4). All conditions used DMEM without pyruvate, with the exception of AA5, which used DMEM with 1 mM pyruvate. **b,** representative Western blots showing expression of vinculin (loading control), GOT1, GOT2, and GFP (jAspSnFR3) in jAspSnFR3-expressing (jAsp), GOT1/2 DKO cells (DKO), and jAspSnFR3-expressing GOT1/2 DKO (jAsp DKO) 143B and H1299 cells. **c-d**, average proliferation rates of 143B (**c**) or H1299 (**d**) GOT1/2 DKO cells (determined using a conventional, endpoint proliferation assay) in DMEM with the indicated initial aspartate concentrations (n=3). **e-f**, relative aspartate levels (green line, left y-axis) and absolute proliferation rates (purple points, right y-axis) of H1299 cells treated with DMSO in DMEM without pyruvate (e) or with 1 mM pyruvate (f) (n=4). **g**, relative aspartate levels and absolute proliferation rates of H1299 cells treated with 75 nM rotenone in DMEM without pyruvate (n=4). **h**, relative aspartate levels and absolute proliferation rates of GOT1/2 DKO H1299 cells after switching from 40 mM aspartate into 2 mM aspartate in DMEM without pyruvate (n=4). **i**, relative aspartate levels and absolute proliferation rates of H1299 cells in DMEM with 1 mM pyruvate treated with 1.5 μM Atpenin A5 (AA5) (n=4). **j**, comparison of the degree of aspartate rebound measured in the experiments in panels g-i (n=4). Data represented as mean +/− S.D. Statistical significance determined using an ordinary one-way ANOVA with uncorrected Fisher’s LSD and a single pooled variance.

**Extended Data Figure 3.**
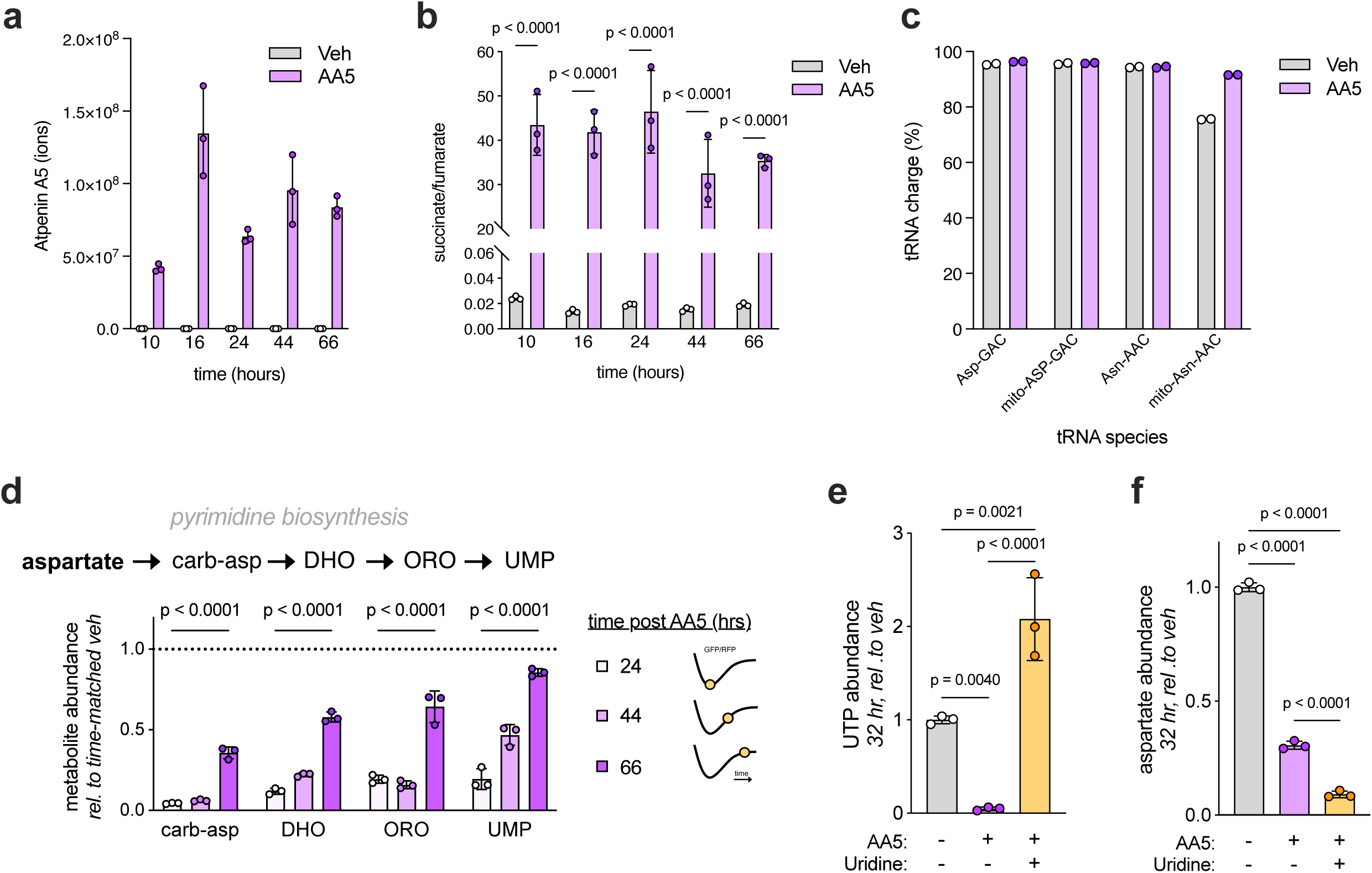
AA5 detection and quantification of aspartate fates in AA5-treated cells. **a**, ion counts of Atpenin A5 (AA5) measured using LC-MS of 143B cells at several timepoints following treatment with vehicle control or 5 μM AA5 (n=3). **b**, succinate/fumarate ratios measured using LC-MS of 143B cells at several timepoints following treatment with 5 μM AA5 or vehicle control (n=3). **c**, absolute charge of four cytosolic and mitochondrial aspartate and asparagine tRNA species measured in 143B cells 30 hours after treatment with 5 μM AA5 or vehicle control (n=2). **d,** relative abundances of carbamoyl-aspartate (carb-asp), dihydroorotate (DHO), orotate (ORO), and uridine monophosphate (UMP) measured using LC-MS of 143B cells at several timepoints following treatment with 5 μM AA5 (n=3). Each abundance is normalized to the corresponding time-matched vehicle-treated control; no change compared to vehicle is represented by the dotted line at y=1. Cartoons illustrate the approximate location of each timepoint on the prototypical GFP/RFP rebound curve in AA5-treated cells. **e,** relative uridine triphosphate (UTP) levels measured using LC-MS of 143B cells at 32 hours post-treatment with vehicle control, 5 μM AA5, or 5 μM AA5 and 200 μΜ uridine (n=3). **f,** relative aspartate levels measured using LC-MS of 143B cells at 32 hours post-treatment with vehicle control, 5 μM AA5, or 5 μM AA5 and 200 μΜ uridine (n=3). Unless otherwise noted, experiments were conducted in DMEM with 1 mM pyruvate. Data represented as mean +/− S.D. Statistical significance determined using an ordinary two-way ANOVA with uncorrected Fisher’s LSD and a single pooled variance (panels b, e, and f) or ordinary two-way ANOVA with a single pooled variance and Tukey’s multiple comparisons correction (panel d).

**Extended Data Figure 4.**
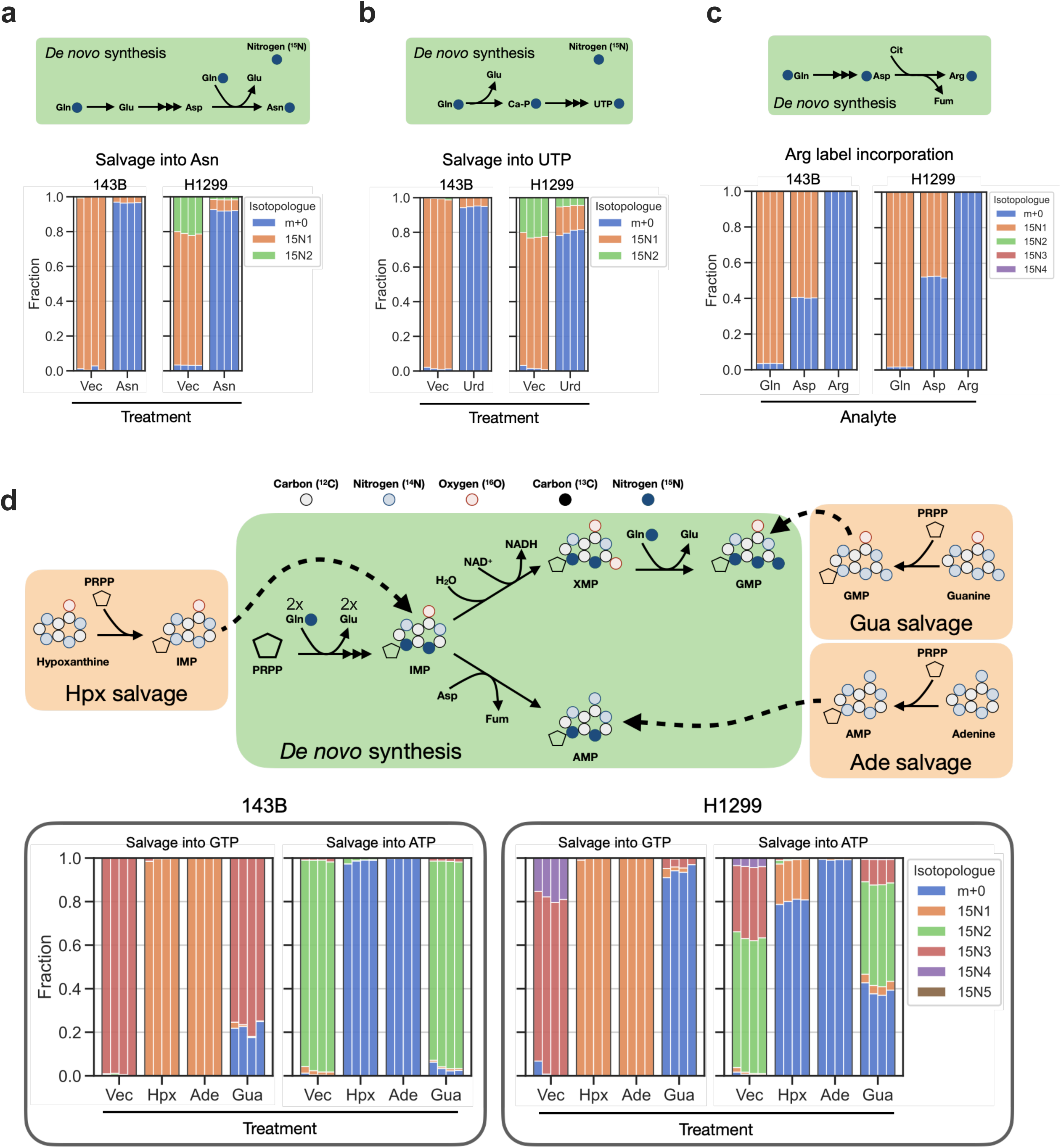
Identification of nutrient conditions to effectively salvage aspartate fates. **a**, diagram demonstrating glutamine (Gln) amide-^15^N label incorporation into asparagine (Asn) and Gln amide-^15^N label incorporation into Asn at steady-state in 143B and H1299 cells grown in DMEM supplemented with vehicle (Vec), or 500 µM Asn (n=4, replicates plotted as individual bars). **b**, diagram demonstrating Gln amide-^15^N label incorporation into uridine triphosphate (UTP) and Gln amide-^15^N label incorporation into UTP at steady-state in 143B and H1299 cells grown in DMEM supplemented with vehicle (Vec), or 200 µM uridine (Urd) (n=4, replicates plotted as individual bars). **c**, diagram demonstrating Gln alpha-^15^N label incorporation into aspartate and subsequently arginine (Arg) and isotopologue distributions for Gln, Asp, and Arg at steady state in 143B and H1299 cells grown in DMEM with no salvageable metabolites added (n=4, replicates plotted as individual bars). **d**, diagram demonstrating Gln amide-^15^N label incorporation into purines and the resulting changes following salvage of hypoxanthine (Hpx), adenine (Ade) or guanine (Gua). AMP/ATP and GMP/GTP are either derived from *de novo* synthesis or salvage. AMP can be salvaged from adenine directly or through hypoxanthine/IMP as a product of adenine deamination. Bottom isotopologue distributions show Gln amide-^15^N label incorporation into GTP and ATP at steady-state in 143B and H1299 cells grown in DMEM supplemented with vehicle (Vec) or 100 µM Hpx, Ade or Gua (n=4, replicates plotted as individual bars).

**Extended Data Figure 5.**
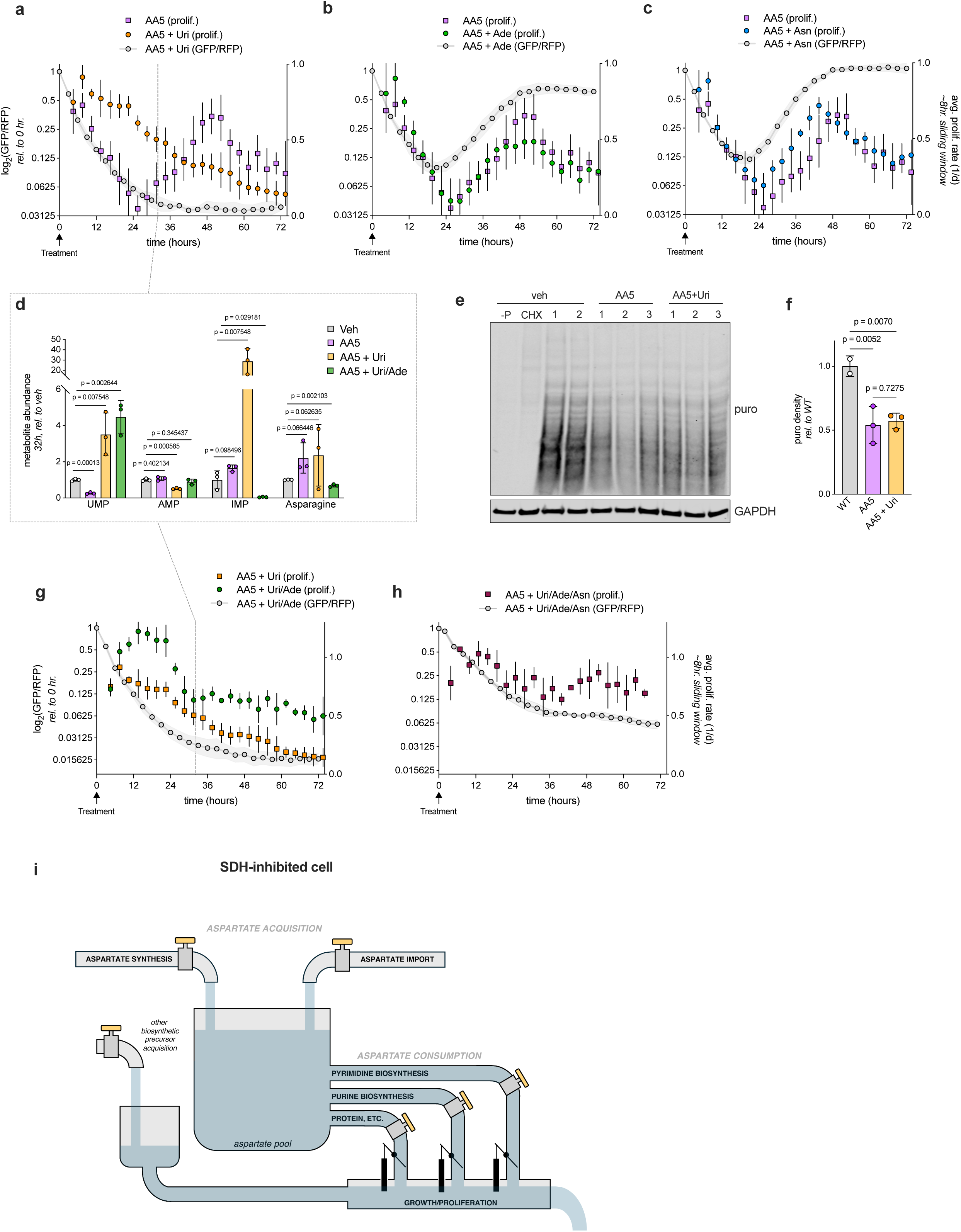
Distributed and hierarchical metabolic growth limitations upon SDH inhibition. **a**, relative aspartate levels (grey line, left y-axis) and absolute proliferation rates (individual points, right y-axis) of 143B sensor cells treated with 5 μM AA5 with or without 200 μΜ uridine (Uri) (n=4). Dotted line marks timepoint corresponding to data in panel d. **b**, relative aspartate levels and absolute proliferation rates of 143B sensor cells treated with 5 μM AA5 with or without 100 μΜ adenine (Ade) (n=4). **c**, relative aspartate levels and absolute proliferation rates of 143B sensor cells treated with 5 μM AA5 with or without 500 μΜ asparagine (Asn) (n=4). **d**, relative levels of UMP, AMP, IMP, and asparagine (Asn) measured using LC-MS in 143B cells 32 hours after treatment with vehicle control, 5 μM AA5, 5 μM AA5 with 200 μΜ uridine (Uri), or 5 μM AA5 with 200 μΜ uridine (Uri) and 100 μM adenine (Ade) (n=3). **e**, 24-hour puromycin incorporation assay using 143B cells treated with vehicle control (veh), 5 μM AA5, or 5 μM AA5 with 200 μΜ uridine (Uri). –P denotes a no-puromycin control, and CHX denotes cells treated with 1 μg/mL of the translation inhibitor cycloheximide. GAPDH serves as a loading control (n=2 for veh, n=3 for AA5 +/− Uri). **f**, quantification of relative puromycin lane densities in panel e (n=2-3). **g**, relative aspartate levels and absolute proliferation rates of 143B sensor cells treated with 5 μM AA5 and 200 μM uridine (Uri, reproduced from panel a) or 200 μM uridine and 100 μΜ adenine (Uri/Ade). Dotted line marks timepoint corresponding to data in panel d (n=4). **h**, relative aspartate levels and absolute proliferation rates of 143B sensor cells treated with 5 μM AA5, 200 μM uridine, 100 μΜ adenine, and 500 μM asparagine (Uri/Ade/Asn) (n=4). **i**, schematic illustrating a conceptual model of aspartate metabolism in SDH-inhibited cells. The aspartate pool is filled by aspartate acquisition (consisting of aspartate synthesis and import) and drained as aspartate is consumed into pyrimidines, purines, and protein, which have an obligatory ‘order.’ Upon constraining aspartate acquisition, aspartate levels fall until they no longer fulfill pyrimidine demands, which closes the gate to stop cell growth/proliferation regardless of the flow through the other aspartate fate/biosynthesis precursor pipes. Note that complementing any single aspartate fate pipe (for example, by supplementing uridine) permits continued aspartate consumption until levels fall to limit the next lowest fate pipe. Unless otherwise noted, experiments were conducted in DMEM with 1 mM pyruvate. Data represented as mean +/− S.D. Statistical significance determined using multiple unpaired t-tests and the two-stage step-up method of Benjamini, Krieger, and Yekutieli to account for multiple comparisons (panel d) or an ordinary one-way ANOVA with uncorrected Fisher’s LSD and a single pooled variance (panel f).

**Extended Data Figure 6.**
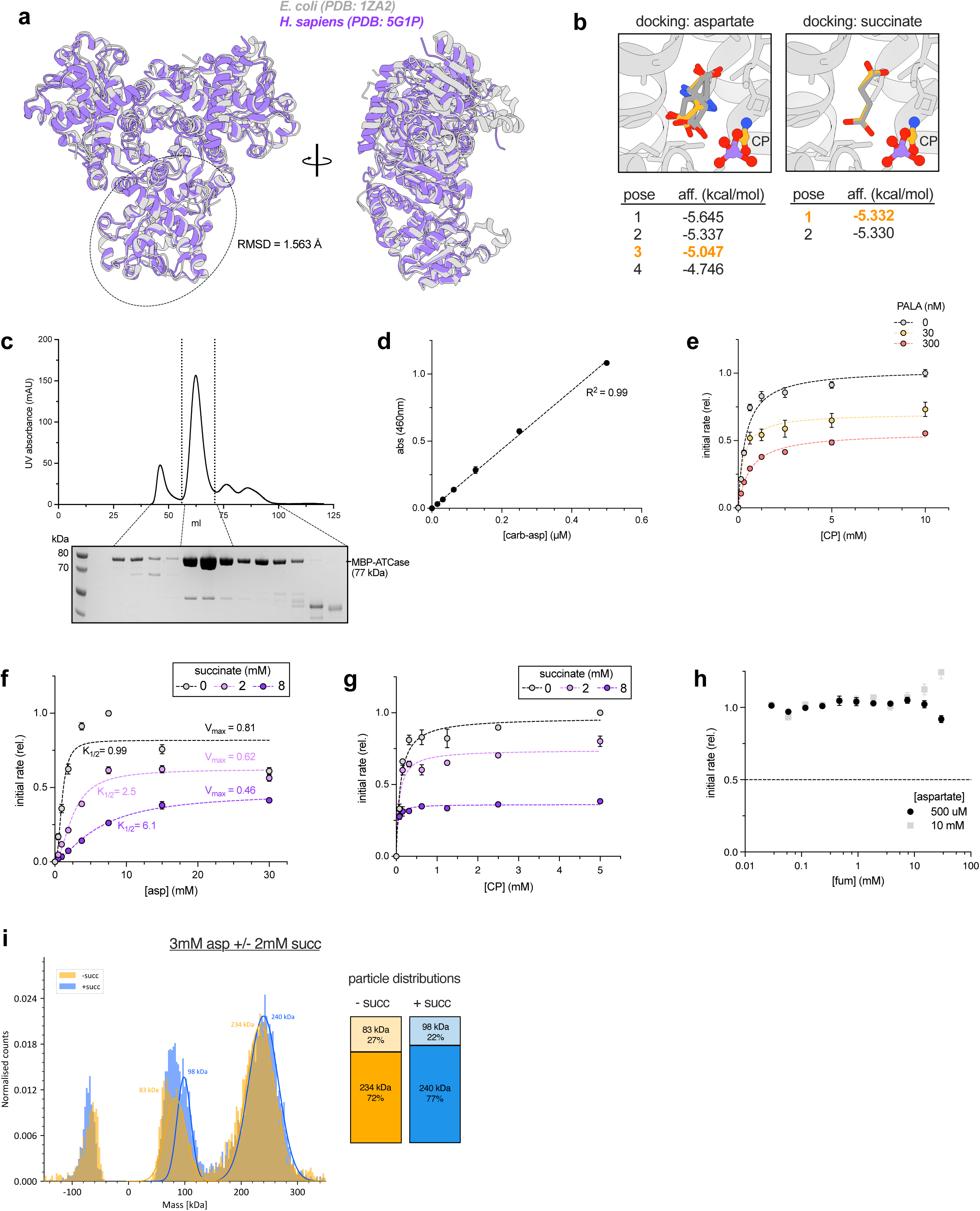
related to Figure 3. **a**, alignment of previously solved crystal structures of *E. coli* (PDB: 1ZA2) and human (PDB: 5G1P) ATCase catalytic trimers. Root mean square deviation (RMSD) reflects structural similarity between the circled monomers. **b**, predicted binding poses and affinities for the top four-scoring aspartate poses and the top two-scoring succinate poses using molecular docking analyses. Poses depicted in Figure 3 are highlighted in yellow. Ligands are colored by atom: yellow/grey = carbon, red = oxygen, blue = nitrogen, purple = phosphorus **c**, UV absorbance trace from size-exclusion chromatography (SEC) of MBP-ATCase affinity chromatography elute, with accompanying Coomassie-stained protein gel measuring MBP-ATCase abundance in the fraction ranges indicated by the dotted lines. Vertical dotted lines indicate fractions taken for downstream analyses. **d**, representative calibration curve for ATCase activity assay plotting absorbance as a function of carbamoyl-aspartate (carb-asp) concentration and fit with linear regression using Prism (n=3 technical replicates). **e**, initial enzymatic rates for purified MBP-ATCase incubated with 5 mM aspartate and the indicated concentrations of CP and PALA (n=3 technical replicates). Data are fit to a Michaelis-Menten model using Prism and rates are normalized to the 10 mM CP, 0 nM PALA condition. **f**, data from Figure 3j fit with cooperative sigmoidal curves using Prism. V_max_ and K_1/2_ values are shown next to the respective conditions. **g**, initial enzymatic rates for purified MBP-ATCase incubated with 10 mM aspartate and the indicated concentrations of CP and succinate (n=3 technical replicates). Data are fit to a Michaelis-Menten model using Prism and rates are normalized to the 10 mM CP, 0 nM succinate condition. **h**, initial enzymatic rates for purified MBP-ATCase incubated with 10 mM CP and the indicated concentrations of aspartate and fumarate (fum) (n=3 technical replicates). Rates are normalized to the 0 mM fumarate condition for each respective aspartate concentration. **i**, particle size distributions obtained using mass photometry on isolated MBP-ATCase incubated with 5 mM CP, 3 mM aspartate, and with or without 2 mM succinate (+/− succ). Predicted molecular weights are labeled above the corresponding Gaussian fits. **j**, quantification of particle distributions in (i), excluding peaks which are mirrored on either side of 0 kDa, a common buffer artifact seen in MP experiments.

**Extended Data Figure 7.**
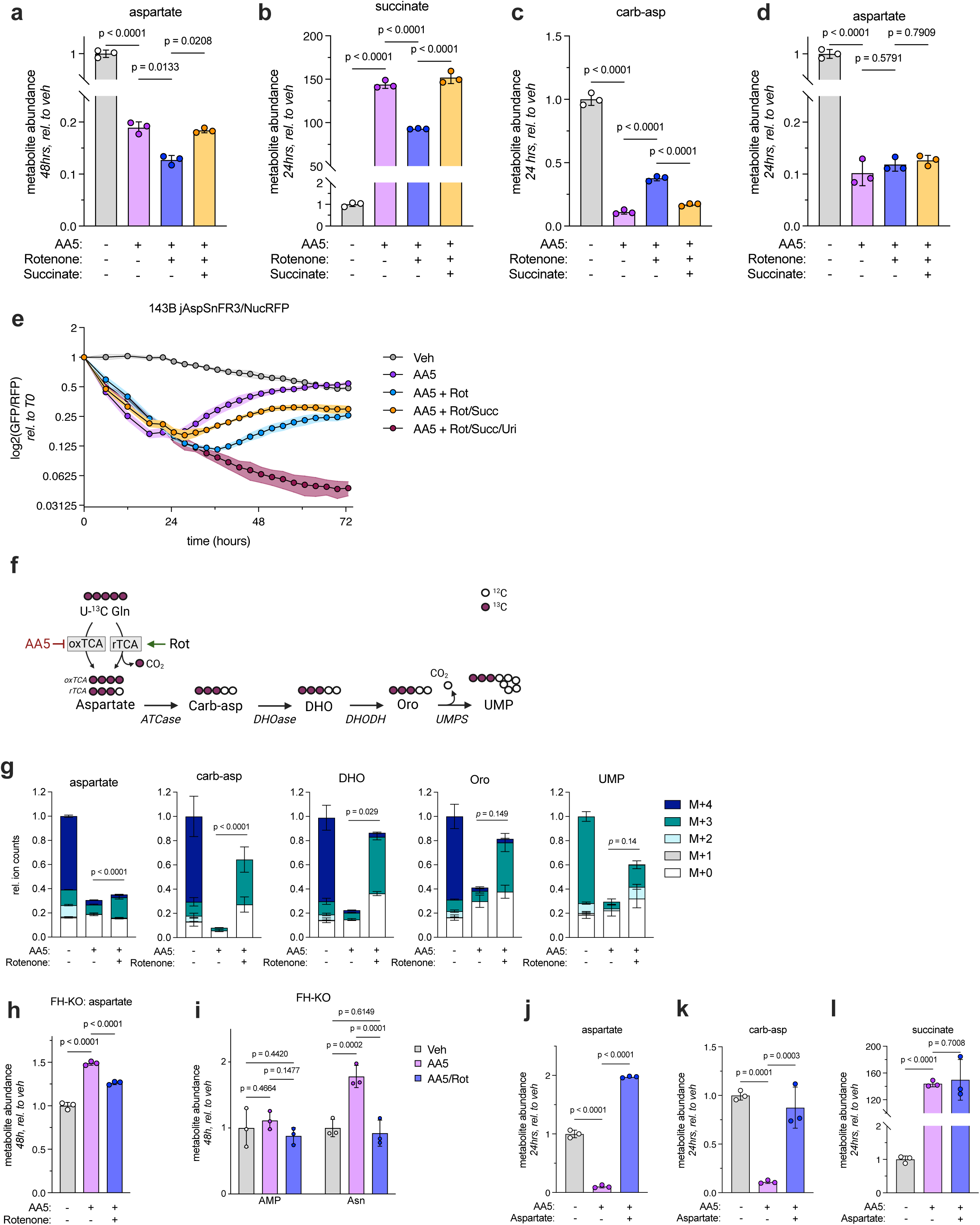
related to Figure 4. **a**, relative aspartate abundances measured by LC-MS on 143B cells 48 hours after treatment with vehicle control or the indicated combinations of 5 μM AA5, 50 nM rotenone, and 10 mM succinate (n=3). **b-d**, relative succinate (b), carbamoyl-aspartate (c), and aspartate (d) abundances measured by LC-MS on 143B cells 24 hours after treatment with vehicle control or the indicated combinations of 5 μM AA5, 50 nM rotenone, and 10 mM succinate (n=3). **e**, relative aspartate levels of 143B sensor cells treated for 72 hours with the indicated combinations of 5 μM AA5, 50 nM rotenone (Rot), 10 mM succinate (Succ), and 200 μΜ uridine (Uri) (n=3). **f**, schematic illustrating U-^13^C glutamine labeling into aspartate and downstream pyrimidine synthesis intermediates via reductive TCA cycling (rTCA), which is stimulated by rotenone co-treatment^7^. **g**, isotopologue distributions of aspartate, carbamoyl-aspartate (carb-asp), dihydroorotate (DHO), orotate (ORO), and uridine monophosphate (UMP) after 30 hours of U-^13^C glutamine tracing in 143B cells treated with vehicle control, 5 μM AA5, or 5 μM AA5 and 50 nM rotenone (n=3). **h**, relative aspartate abundances measured by LC-MS on FH-KO 143B cells 48 hours after treatment with vehicle control, 5 μM AA5, or 5 μM AA5 and 50 nM rotenone (n=3). **i**, relative AMP and asparagine (Asn) abundances measured by LC-MS on FH-KO 143B cells 48 hours after treatment with vehicle control, 5 μM AA5, or 5 μM AA5 and 50 nM rotenone (Rot) (n=3). **j-l**, relative aspartate (j), carbamoyl-aspartate (carb-asp) (k), and succinate (l) abundances measured by LC-MS on 143B cells 24 hours after treatment with vehicle control, 5 μM AA5, or 5 μΜ ΑΑ5 and 20 mM aspartate (n=3). Unless otherwise noted, experiments were conducted in DMEM with 1 mM pyruvate. Data represented as mean +/− S.D. Statistical significance determined using an ordinary two-way ANOVA with uncorrected Fisher’s LSD and a single pooled variance, except panel h, which used an ordinary two-way ANOVA with main effects only, uncorrected Fisher’s LSD and a single pooled variance. For panel h, p values denote the results of statistical testing on the M+3 fraction of each metabolite. ATCase, aspartate transcarbamylase; DHOase, dihydroorotase; DHODH, dihydroorotate dehydrogenase; UMPS, UMP synthase.

**Extended Data Figure 8.**
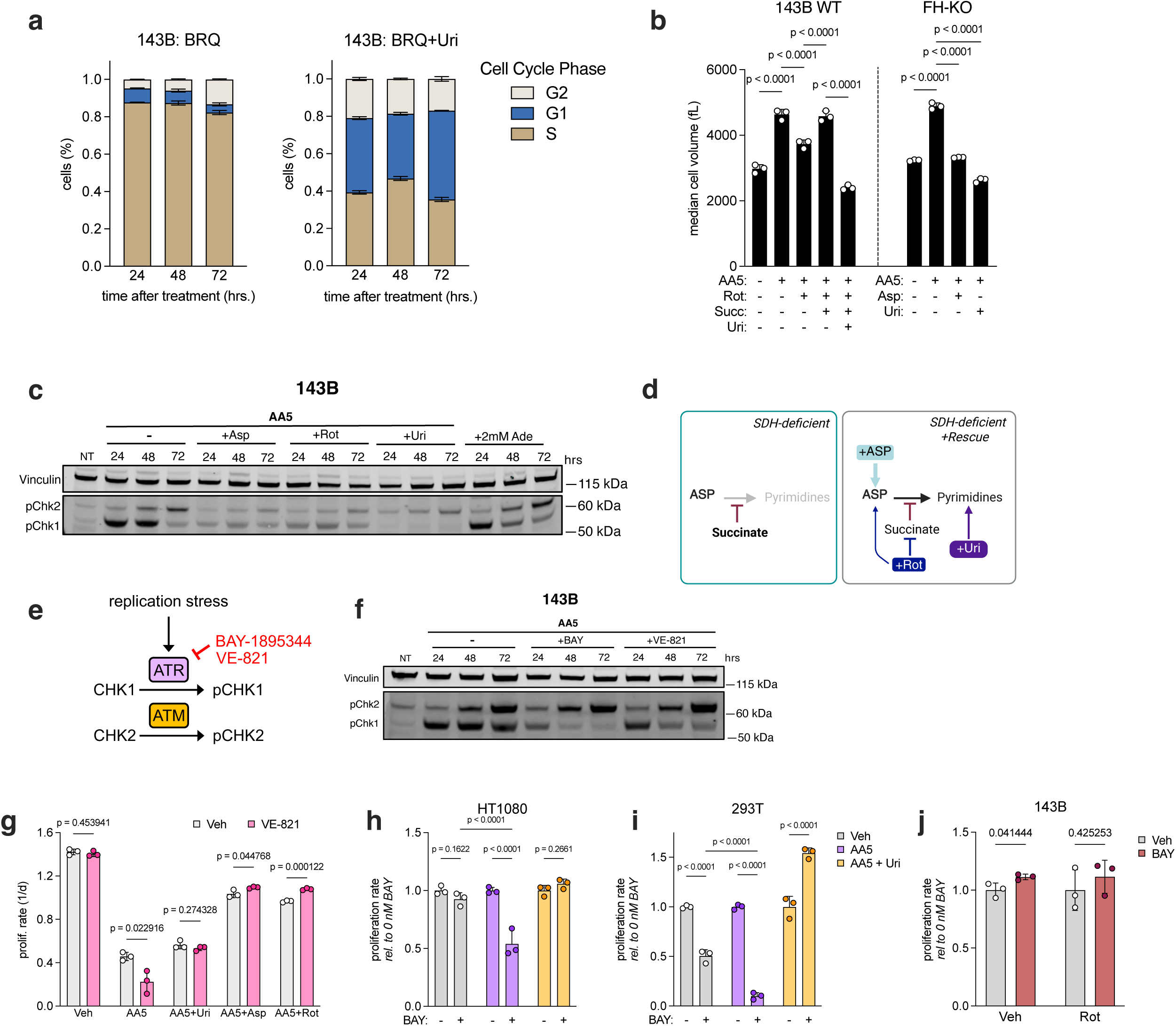
related to Figure 5. **a**, representative cell cycle experiment showing the proportion of 143B cells in each cell cycle phase at the indicated timepoints following treatment with 2 μM brequinar (BRQ) with or without 200 μM uridine (Uri) (n=3). **b**, median cell volumes of 143B wild-type (WT) or 143B FH knockout (FH-KO) cells after 72 hours of treatment with the indicated combinations of 5 μM AA5, 50 nM rotenone (Rot), 20 mM succinate (Succ), 200 μM uridine (Uri), and 20 mM aspartate (Asp) (n=3). **c**, representative Western blot demonstrating levels of phosphorylated CHK1 (pCHK1), phosphorylated CHK2 (pCHK2), and vinculin loading control in 143B cells at the indicated timepoints following treatment with control (NT), 2 mM adenine (Ade), 5 μΜ ΑΑ5, or 5 μΜ ΑΑ5 with either 20 mM aspartate (Asp), 50 nM rotenone (Rot), or 200 μM uridine (Uri). **d**, schematic illustrating various metabolic rescues of pyrimidine synthesis in SDH-deficient cells. **e**, identical schematic as figure 5e, with the addition of the ATR inhibitor VE-821 and ATM kinase, which phosphorylates CHK2. **f,** representative Western blot demonstrating levels of phosphorylated CHK1 (pCHK1), phosphorylated CHK2 (pCHK2), and vinculin loading control in 143B cells at the indicated timepoints following treatment with control (NT), 5 μΜ ΑΑ5, or 5 μΜ ΑΑ5 with either 20 nM BAY-1895344 or 0.6 μΜ VE-821. **g**, absolute proliferation rates (measured using a conventional, 72-hour endpoint proliferation assay) of 143B cells treated with vehicle control or 0.6 μM VE-821 and the indicated combinations of vehicle control, 5 μM AA5, 200 μM uridine (Uri), 20 mM aspartate (Asp), and 50 nM rotenone (Rot) (n=3). **h-i**, relative proliferation rates (measured using a conventional, 72-hour endpoint proliferation assay) of HT1080 (h) and 293T (i) cells treated with the indicated combinations of 5 μM AA5 and either 10 nM (h) or 50 nM (i) BAY-1895344 (BAY) (n=3). **j**, relative proliferation rates (measured using a conventional, 72-hour endpoint proliferation assay) of 143B cells treated with the indicated combinations of 50 nM rotenone (Rot) and either vehicle control or 20 nM BAY-1895344 (BAY) in DMEM without pyruvate (n=3). Unless otherwise noted, experiments were conducted in DMEM with 1 mM pyruvate. Data represented as mean +/− S.D. Statistical significance determined using an ordinary one-way ANOVA with Tukey’s multiple comparisons test and a single pooled variance (panel b), ordinary two-way ANOVAs with uncorrected Fisher’s LSD and a single pooled variance (panels h-i) or multiple unpaired t-tests (panels g, j).

**Extended Data Figure 9.**
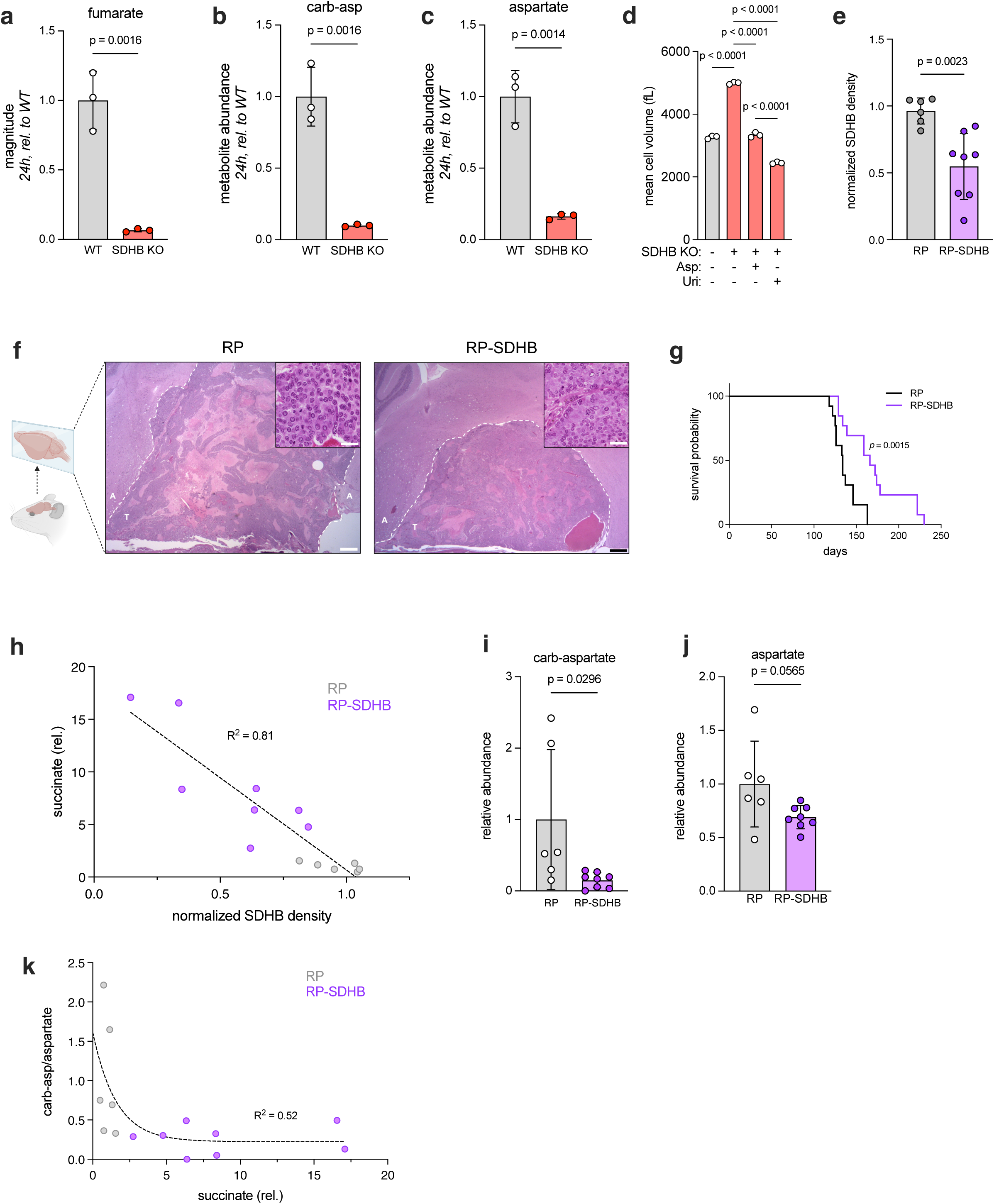
Related to Figure 6. **a-c**, relative fumarate (a), carbamoyl-aspartate (b), and aspartate (c) abundances measured by LC-MS on SDHB-KO 143B cells 24 hours after media change (n=3). **d** mean cell volumes of wild type or SDHB-KO 143B cells 48 hours after treatment with vehicle control, 20 mM aspartate or 200 μM uridine (n=3). **e**, relative SDHB band density in tumor extracts from *Rb*^−/−^, *Tp53*^−/−^, *Sdhb*^−/−^ (RP-SDHB) or *Rb*^−/−^, *Tp53*^−/−^ mice (RP) calculated from the blot in Figure 6i (n=6 RP, 8 RP-SDHB). **f**, representative hematoxylin and eosin (H&E) stained, formalin-fixed tumor sections from one RP and RP-SDHB mouse. Larger images are 2X magnification, insets are 60X. Dotted lines separate tumor (T) from adjacent (A) tissue, 2X scale bar = 600 μm, inset scale bar = 20 μm. **g**, Kaplan-Meier plot demonstrating overall survival of RP-SDHB and RP mice (n=13). **h**, normalized SDHB band density from Figure 6i plotted against relative succinate abundances from Figure 6j on a per-tumor basis. Dotted line and R^2^ value show results of linear regression analysis (n=6 RP, 8 RP-SDHB). **i-j**, relative carbamoyl-aspartate (i) and aspartate (j) abundances measured by LC-MS on pituitary tumor extracts from RP and RP-SDHB mice (n=6 RP, 8 RP-SDHB). **k**, relative succinate abundances from Figure 6j plotted against carbamoyl-aspartate/aspartate ratios from Figure 6k on a per-sample basis. Dotted line and R^2^ value show the results of nonlinear regression analysis fitting a one-phase exponential decay (n=6 RP, 8 RP-SDHB). Unless otherwise noted, experiments were conducted in DMEM with 1 mM pyruvate, data represented as median +/− quartiles (panels h-i) or mean +/− S.D. (all other panels). Statistical significance determined using unpaired t-tests (panels a-c, e, i-j) Mantel-Cox test (panel g) or an ordinary one-way ANOVA with uncorrected Fisher’s LSD and a single pooled variance (panel d).

## References

1. Shamir, M., Bar-On, Y., Phillips, R., and Milo, R. (2016). SnapShot: Timescales in Cell Biology. Cell 164, 1302–1302.e1. 10.1016/j.cell.2016.02.058.

2. Sullivan, L.B., Gui, D.Y., Hosios, A.M., Bush, L.N., Freinkman, E., and Vander Heiden, M.G. (2015). Supporting Aspartate Biosynthesis Is an Essential Function of Respiration in Proliferating Cells. Cell 162, 552–563. 10.1016/j.cell.2015.07.017.

3. Birsoy, K., Wang, T., Chen, W.W., Freinkman, E., Abu-Remaileh, M., and Sabatini, D.M. (2015). An Essential Role of the Mitochondrial Electron Transport Chain in Cell Proliferation Is to Enable Aspartate Synthesis. Cell 162, 540–551. 10.1016/j.cell.2015.07.016.

4. Gui, D.Y., Sullivan, L.B., Luengo, A., Hosios, A.M., Bush, L.N., Gitego, N., Davidson, S.M., Freinkman, E., Thomas, C.J., and Vander Heiden, M.G. (2016). Environment Dictates Dependence on Mitochondrial Complex I for NAD+ and Aspartate Production and Determines Cancer Cell Sensitivity to Metformin. Cell Metabolism 24, 716–727. 10.1016/j.cmet.2016.09.006.

5. Sullivan, L.B., Luengo, A., Danai, L.V., Bush, L.N., Diehl, F.F., Hosios, A.M., Lau, A.N., Elmiligy, S., Malstrom, S., Lewis, C.A., et al. (2018). Aspartate is an endogenous metabolic limitation for tumour growth. Nat. Cell Biol. 20, 782–788. 10.1038/s41556-018-0125-0.

6. Lussey-Lepoutre, C., Hollinshead, K.E.R., Ludwig, C., Menara, M., Morin, A., Castro-Vega, L.-J., Parker, S.J., Janin, M., Martinelli, C., Ottolenghi, C., et al. (2015). Loss of succinate dehydrogenase activity results in dependency on pyruvate carboxylation for cellular anabolism. Nat Commun 6, 1–9. 10.1038/ncomms9784.

7. Hart, M.L., Quon, E., Vigil, A.-L.B., Engstrom, I.A., Newsom, O.J., Davidsen, K., Hoellerbauer, P., Carlisle, S.M., and Sullivan, L.B. (2023). Mitochondrial redox adaptations enable alternative aspartate synthesis in SDH-deficient cells. eLife 12, e78654. 10.7554/eLife.78654.

8. Cardaci, S., Zheng, L., MacKay, G., van den Broek, N.J.F., MacKenzie, E.D., Nixon, C., Stevenson, D., Tumanov, S., Bulusu, V., Kamphorst, J.J., et al. (2015). Pyruvate carboxylation enables growth of SDH-deficient cells by supporting aspartate biosynthesis. Nat Cell Biol 17, 1317–1326. 10.1038/ncb3233.

9. Mick, E., Titov, D.V., Skinner, O.S., Sharma, R., Jourdain, A.A., and Mootha, V.K. (2020). Distinct mitochondrial defects trigger the integrated stress response depending on the metabolic state of the cell. eLife 9, e49178. 10.7554/eLife.49178.

10. Qi, L., Martin-Sandoval, M.S., Merchant, S., Gu, W., Eckhardt, M., Mathews, T.P., Zhao, Z., Agathocleous, M., and Morrison, S.J. (2021). Aspartate availability limits hematopoietic stem cell function during hematopoietic regeneration. Cell Stem Cell 28, 1982–1999.e8. 10.1016/j.stem.2021.07.011.

11. Garcia-Bermudez, J., Baudrier, L., La, K., Zhu, X.G., Fidelin, J., Sviderskiy, V.O., Papagiannakopoulos, T., Molina, H., Snuderl, M., Lewis, C.A., et al. (2018). Aspartate is a limiting metabolite for cancer cell proliferation under hypoxia and in tumours. Nature Cell Biology 20, 775–781. 10.1038/s41556-018-0118-z.

12. Garcia-Bermudez, J., Badgley, M.A., Prasad, S., Baudrier, L., Liu, Y., La, K., Soula, M., Williams, R.T., Yamaguchi, N., Hwang, R.F., et al. (2022). Adaptive stimulation of macropinocytosis overcomes aspartate limitation in cancer cells under hypoxia. Nat Metab 4, 724–738. 10.1038/s42255-022-00583-z.

13. Madala, H.R., Helenius, I.T., Zhou, W., Mills, E., Zhang, Y., Liu, Y., Metelo, A.M., Kelley, M.L., Punganuru, S., Kim, K.B., et al. (2020). Nitrogen Trapping as a Therapeutic Strategy in Tumors with Mitochondrial Dysfunction. Cancer Research 80, 3492–3506. 10.1158/0008-5472.CAN-20-0246.

14. Kerk, S.A., Lin, L., Myers, A.L., Sutton, D.J., Andren, A., Sajjakulnukit, P., Zhang, L., Zhang, Y., Jiménez, J.A., Nelson, B.S., et al. (2022). Metabolic requirement for GOT2 in pancreatic cancer depends on environmental context. eLife 11, e73245. 10.7554/eLife.73245.

15. Teh, M.R., Gudgeon, N., Frost, J.N., Sinclair, L.V., Smith, A.L., Millington, C.L., Kronsteiner, B., Roberts, J., Marzullo, B.P., Murray, H., et al. (2025). Iron deficiency causes aspartate-sensitive dysfunction in CD8+ T cells. Nat Commun 16, 5355. 10.1038/s41467-025-60204-7.

16. Alkan, H.F., Walter, K.E., Luengo, A., Madreiter-Sokolowski, C.T., Stryeck, S., Lau, A.N., Al-Zoughbi, W., Lewis, C.A., Thomas, C.J., Hoefler, G., et al. (2018). Cytosolic Aspartate Availability Determines Cell Survival When Glutamine Is Limiting. Cell Metab 28, 706–720.e6. 10.1016/j.cmet.2018.07.021.

17. Meléndez-Rodríguez, F., Urrutia, A.A., Lorendeau, D., Rinaldi, G., Roche, O., Böğürcü-Seidel, N., Ortega Muelas, M., Mesa-Ciller, C., Turiel, G., Bouthelier, A., et al. (2019). HIF1α Suppresses Tumor Cell Proliferation through Inhibition of Aspartate Biosynthesis. Cell Reports 26, 2257–2265.e4. 10.1016/j.celrep.2019.01.106.

18. Altea-Manzano, P., Vandekeere, A., Edwards-Hicks, J., Roldan, M., Abraham, E., Lleshi, X., Guerrieri, A.N., Berardi, D., Wills, J., Junior, J.M., et al. (2022). Reversal of mitochondrial malate dehydrogenase 2 enables anaplerosis via redox rescue in respiration-deficient cells. Mol Cell 82, 4537–4547.e7. 10.1016/j.molcel.2022.10.005.

19. Cheng, C.-T., Qi, Y., Wang, Y.-C., Chi, K.K., Chung, Y., Ouyang, C., Chen, Y.-R., Oh, M.E., Sheng, X., Tang, Y., et al. (2018). Arginine starvation kills tumor cells through aspartate exhaustion and mitochondrial dysfunction. Commun Biol 1, 1–15. 10.1038/s42003-018-0178-4.

20. Oberkersch, R.E., Pontarin, G., Astone, M., Spizzotin, M., Arslanbaeva, L., Tosi, G., Panieri, E., Ricciardi, S., Allega, M.F., Brossa, A., et al. (2022). Aspartate metabolism in endothelial cells activates the mTORC1 pathway to initiate translation during angiogenesis. Developmental Cell 57, 1241–1256.e8. 10.1016/j.devcel.2022.04.018.

21. Davidsen, K., Marvin, J.S., Aggarwal, A., Brown, T.A., and Sullivan, L.B. (2023). An engineered biosensor enables dynamic aspartate measurements in living cells. eLife 12. 10.7554/eLife.90024.

22. Marvin, J.S., Borghuis, B.G., Tian, L., Cichon, J., Harnett, M.T., Akerboom, J., Gordus, A., Renninger, S.L., Chen, T.-W., Bargmann, C.I., et al. (2013). An optimized fluorescent probe for visualizing glutamate neurotransmission. Nat Methods 10, 162–170. 10.1038/nmeth.2333.

23. Lendvai, N., Pawlosky, R., Bullova, P., Eisenhofer, G., Patocs, A., Veech, R.L., and Pacak, K. (2014). Succinate-to-fumarate ratio as a new metabolic marker to detect the presence of SDHB/D-related paraganglioma: initial experimental and ex vivo findings. Endocrinology 155, 27–32. 10.1210/en.2013-1549.

24. Davidsen, K., and Sullivan, L.B. (2024). A robust method for measuring aminoacylation through tRNA-Seq. eLife 12, RP91554. 10.7554/eLife.91554.

25. Rabinovich, S., Adler, L., Yizhak, K., Sarver, A., Silberman, A., Agron, S., Stettner, N., Sun, Q., Brandis, A., Helbling, D., et al. (2015). Diversion of aspartate in ASS1-deficient tumours fosters de novo pyrimidine synthesis. Nature 527, 379–383. 10.1038/nature15529.

26. de Kant, E., Pinedo, H.M., Laurensse, E., and Peters, G.J. (1989). The relation between inhibition of cell growth and of dihydroorotic acid dehydrogenase by brequinar sodium. Cancer Lett 46, 123–127. 10.1016/0304-3835(89)90019-0.

27. Dalla Pozza, E., Dando, I., Pacchiana, R., Liboi, E., Scupoli, M.T., Donadelli, M., and Palmieri, M. (2019). Regulation of succinate dehydrogenase and role of succinate in cancer. Semin. Cell Dev. Biol. 10.1016/j.semcdb.2019.04.013.

28. Bennett, B.D., Yuan, J., Kimball, E.H., and Rabinowitz, J.D. (2008). Absolute quantitation of intracellular metabolite concentrations by an isotope ratio-based approach. Nat Protoc 3, 1299–1311. 10.1038/nprot.2008.107.

29. Nengroo, M.A., Klein, A.T., Carr, H.S., Vidal-Cruchez, O., Sahu, U., McGrail, D.J., Sahni, N., Thompson, N.B., Faull, P.A., Gao, P., et al. (2025). Accumulation of succinate suppresses de novo purine synthesis through succinylation-mediated control of the mitochondrial folate cycle. Mol Cell 85, 4215–4228.e9. 10.1016/j.molcel.2025.10.002.

30. Sulkowski, P.L., Sundaram, R.K., Oeck, S., Corso, C.D., Liu, Y., Noorbakhsh, S., Niger, M., Boeke, M., Ueno, D., Kalathil, A.N., et al. (2018). Krebs Cycle-Deficient Hereditary Cancer Syndromes are Defined by Homologous Recombination DNA Repair Defects. Nat Genet 50, 1086–1092. 10.1038/s41588-018-0170-4.

31. Richter, S., Peitzsch, M., Rapizzi, E., Lenders, J.W., Qin, N., de Cubas, A.A., Schiavi, F., Rao, J.U., Beuschlein, F., Quinkler, M., et al. (2014). Krebs Cycle Metabolite Profiling for Identification and Stratification of Pheochromocytomas/Paragangliomas due to Succinate Dehydrogenase Deficiency. J Clin Endocrinol Metab 99, 3903–3911. 10.1210/jc.2014-2151.

32. Letouzé, E., Martinelli, C., Loriot, C., Burnichon, N., Abermil, N., Ottolenghi, C., Janin, M., Menara, M., Nguyen, A.T., Benit, P., et al. (2013). SDH Mutations Establish a Hypermethylator Phenotype in Paraganglioma. Cancer Cell 23, 739–752. 10.1016/j.ccr.2013.04.018.

33. Porter, R.W., Modebe, M.O., and Stark, G.R. (1969). Aspartate Transcarbamylase: Kinetic studies of the catalytic subunit. Journal of Biological Chemistry 244, 1846–1859. 10.1016/S0021-9258(18)91759-X.

34. Collins, K.D., and Stark, G.R. (1971). Aspartate Transcarbamylase: INTERACTION WITH THE TRANSITION STATE ANALOGUE N-(PHOSPHONACETYL)-l-ASPARTATE. Journal of Biological Chemistry 246, 6599–6605. 10.1016/S0021-9258(19)34156-0.

35. Foote, J., Lauritzen, A.M., and Lipscomb, W.N. (1985). Substrate specificity of aspartate transcarbamylase. Interaction of the enzyme with analogs of aspartate and succinate. J Biol Chem 260, 9624–9629.

36. Ruiz-Ramos, A., Velázquez-Campoy, A., Grande-García, A., Moreno-Morcillo, M., and Ramón-Maiques, S. (2016). Structure and Functional Characterization of Human Aspartate Transcarbamoylase, the Target of the Anti-tumoral Drug PALA. Structure 24, 1081–1094. 10.1016/j.str.2016.05.001.

37. Del Caño-Ochoa, F., Moreno-Morcillo, M., and Ramón-Maiques, S. (2019). CAD, A Multienzymatic Protein at the Head of de Novo Pyrimidine Biosynthesis. In Macromolecular Protein Complexes II: Structure and Function Subcellular Biochemistry., J. R. Harris and J. Marles-Wright, eds. (Springer International Publishing), pp. 505–538. 10.1007/978-3-030-28151-9_17.

38. Lipscomb, W.N., and Kantrowitz, E.R. (2012). Structure and Mechanisms of Escherichia coli Aspartate Transcarbamoylase. Acc. Chem. Res. 45, 444–453. 10.1021/ar200166p.

39. Else, A.J., and Hervé, G. (1990). A microtiter plate assay for aspartate transcarbamylase. Anal Biochem 186, 219–221. 10.1016/0003-2697(90)90069-l.

40. Swyryd, E.A., Seaver, S.S., and Stark, G.R. (1974). N-(phosphonacetyl)-L-aspartate, a potent transition state analog inhibitor of aspartate transcarbamylase, blocks proliferation of mammalian cells in culture. J Biol Chem 249, 6945–6950.

41. Del Caño-Ochoa, F., and Ramón-Maiques, S. (2021). Deciphering CAD: Structure and function of a mega-enzymatic pyrimidine factory in health and disease. Protein Sci 30, 1995–2008. 10.1002/pro.4158.

42. Lorendeau, D., Rinaldi, G., Boon, R., Spincemaille, P., Metzger, K., Jäger, C., Christen, S., Dong, X., Kuenen, S., Voordeckers, K., et al. (2017). Dual loss of succinate dehydrogenase (SDH) and complex I activity is necessary to recapitulate the metabolic phenotype of SDH mutant tumors. Metabolic Engineering 43, 187–197. 10.1016/j.ymben.2016.11.005.

43. Yoo, A., Tang, C., Zucker, M., Fitzgerald, K., DiNatale, R.G., Rappold, P.M., Weiss, K., Freeman, B., Lee, C.-H., Schultz, N., et al. (2022). Genomic and Metabolic Hallmarks of SDH– and FH-deficient Renal Cell Carcinomas. Eur Urol Focus 8, 1278–1288. 10.1016/j.euf.2021.12.002.

44. Blackford, A.N., and Jackson, S.P. (2017). ATM, ATR, and DNA-PK: The Trinity at the Heart of the DNA Damage Response. Molecular Cell 66, 801–817. 10.1016/j.molcel.2017.05.015.

45. Gaillard, H., García-Muse, T., and Aguilera, A. (2015). Replication stress and cancer. Nat Rev Cancer 15, 276–289. 10.1038/nrc3916.

46. Do, B.T., Hsu, P.P., Vermeulen, S.Y., Wang, Z., Hirz, T., Abbott, K.L., Aziz, N., Replogle, J.M., Bjelosevic, S., Paolino, J., et al. (2024). Nucleotide depletion promotes cell fate transitions by inducing DNA replication stress. Dev Cell 59, 2203–2221.e15. 10.1016/j.devcel.2024.05.010.

47. Diehl, F.F., Miettinen, T.P., Elbashir, R., Nabel, C.S., Darnell, A.M., Do, B.T., Manalis, S.R., Lewis, C.A., and Vander Heiden, M.G. (2022). Nucleotide imbalance decouples cell growth from cell proliferation. Nat Cell Biol 24, 1252–1264. 10.1038/s41556-022-00965-1.

48. Bartek, J., Lukas, C., and Lukas, J. (2004). Checking on DNA damage in S phase. Nat Rev Mol Cell Biol 5, 792–804. 10.1038/nrm1493.

49. Smith, J., Tho, L.M., Xu, N., and Gillespie, D.A. (2010). The ATM-Chk2 and ATR-Chk1 pathways in DNA damage signaling and cancer. Adv Cancer Res 108, 73–112. 10.1016/B978-0-12-380888-2.00003-0.

50. Maréchal, A., and Zou, L. (2013). DNA damage sensing by the ATM and ATR kinases. Cold Spring Harb Perspect Biol 5, a012716. 10.1101/cshperspect.a012716.

51. Wengner, A.M., Siemeister, G., Lücking, U., Lefranc, J., Wortmann, L., Lienau, P., Bader, B., Bömer, U., Moosmayer, D., Eberspächer, U., et al. (2020). The Novel ATR Inhibitor BAY 1895344 Is Efficacious as Monotherapy and Combined with DNA Damage-Inducing or Repair-Compromising Therapies in Preclinical Cancer Models. Mol Cancer Ther 19, 26–38. 10.1158/1535-7163.MCT-19-0019.

52. Prevo, R., Fokas, E., Reaper, P.M., Charlton, P.A., Pollard, J.R., McKenna, W.G., Muschel, R.J., and Brunner, T.B. (2012). The novel ATR inhibitor VE-821 increases sensitivity of pancreatic cancer cells to radiation and chemotherapy. Cancer Biol Ther 13, 1072–1081. 10.4161/cbt.21093.

53. Gill, A.J., Toon, C.W., Clarkson, A., Sioson, L., Chou, A., Winship, I., Robinson, B.G., Benn, D.E., Clifton-Bligh, R.J., and Dwight, T. (2014). Succinate Dehydrogenase Deficiency Is Rare in Pituitary Adenomas. Am J Surg Pathol 38, 560–566. 10.1097/PAS.0000000000000149.

54. Astuti, D., Latif, F., Dallol, A., Dahia, P.L.M., Douglas, F., George, E., Sköldberg, F., Husebye, E.S., Eng, C., and Maher, E.R. (2001). Gene Mutations in the Succinate Dehydrogenase Subunit SDHB Cause Susceptibility to Familial Pheochromocytoma and to Familial Paraganglioma. The American Journal of Human Genetics 69, 49–54. 10.1086/321282.

55. Baysal, B.E., Ferrell, R.E., Willett-Brozick, J.E., Lawrence, E.C., Myssiorek, D., Bosch, A., Mey, A. van der, Taschner, P.E.M., Rubinstein, W.S., Myers, E.N., et al. (2000). Mutations in SDHD, a Mitochondrial Complex II Gene, in Hereditary Paraganglioma. Science 287, 848–851. 10.1126/science.287.5454.848.

56. Niemann, S., and Müller, U. (2000). Mutations in SDHC cause autosomal dominant paraganglioma, type 3. Nat. Genet. 26, 268–270. 10.1038/81551.

57. Janeway, K.A., Kim, S.Y., Lodish, M., Nosé, V., Rustin, P., Gaal, J., Dahia, P.L.M., Liegl, B., Ball, E.R., Raygada, M., et al. (2011). Defects in succinate dehydrogenase in gastrointestinal stromal tumors lacking KIT and PDGFRA mutations. Proc. Natl. Acad. Sci. U.S.A. 108, 314–318. 10.1073/pnas.1009199108.

58. Xekouki, P., Szarek, E., Bullova, P., Giubellino, A., Quezado, M., Mastroyannis, S.A., Mastorakos, P., Wassif, C.A., Raygada, M., Rentia, N., et al. (2015). Pituitary adenoma with paraganglioma/pheochromocytoma (3PAs) and succinate dehydrogenase defects in humans and mice. J Clin Endocrinol Metab 100, E710–719. 10.1210/jc.2014-4297.

59. Bardella, C., Pollard, P.J., and Tomlinson, I. (2011). SDH mutations in cancer. Biochim. Biophys. Acta 1807, 1432–1443. 10.1016/j.bbabio.2011.07.003.

60. Armstrong, N., Storey, C.M., Noll, S.E., Margulis, K., Soe, M.H., Xu, H., Yeh, B., Fishbein, L., Kebebew, E., Howitt, B.E., et al. (2022). SDHB knockout and succinate accumulation are insufficient for tumorigenesis but dual SDHB/NF1 loss yields SDHx-like pheochromocytomas. Cell Reports 38, 110453. 10.1016/j.celrep.2022.110453.

61. Jang, C., Chen, L., and Rabinowitz, J.D. (2018). Metabolomics and Isotope Tracing. Cell 173, 822–837. 10.1016/j.cell.2018.03.055.

62. Eijkelenkamp, K., Osinga, T.E., Links, T.P., and Horst-Schrivers, A.N.A. van der Clinical implications of the oncometabolite succinate in SDHx-mutation carriers. Clinical Genetics 0. 10.1111/cge.13553.

63. Chouchani, E.T., Pell, V.R., Gaude, E., Aksentijević, D., Sundier, S.Y., Robb, E.L., Logan, A., Nadtochiy, S.M., Ord, E.N.J., Smith, A.C., et al. (2014). Ischaemic accumulation of succinate controls reperfusion injury through mitochondrial ROS. Nature 515, 431–435. 10.1038/nature13909.

64. Tannahill, G.M., Curtis, A.M., Adamik, J., Palsson-McDermott, E.M., McGettrick, A.F., Goel, G., Frezza, C., Bernard, N.J., Kelly, B., Foley, N.H., et al. (2013). Succinate is an inflammatory signal that induces IL-1β through HIF-1α. Nature 496, 238–242. 10.1038/nature11986.

65. Sullivan, L.B., Gui, D.Y., and Vander Heiden, M.G. (2016). Altered metabolite levels in cancer: implications for tumour biology and cancer therapy. Nat. Rev. Cancer 16, 680–693. 10.1038/nrc.2016.85.

66. Mills, E.L., Kelly, B., Logan, A., Costa, A.S.H., Varma, M., Bryant, C.E., Tourlomousis, P., Däbritz, J.H.M., Gottlieb, E., Latorre, I., et al. (2016). Succinate Dehydrogenase Supports Metabolic Repurposing of Mitochondria to Drive Inflammatory Macrophages. Cell 167, 457–470.e13. 10.1016/j.cell.2016.08.064.

67. Keiran, N., Ceperuelo-Mallafré, V., Calvo, E., Hernández-Alvarez, M.I., Ejarque, M., Núñez-Roa, C., Horrillo, D., Maymó-Masip, E., Rodríguez, M.M., Fradera, R., et al. (2019). SUCNR1 controls an anti-inflammatory program in macrophages to regulate the metabolic response to obesity. Nat Immunol 20, 581–592. 10.1038/s41590-019-0372-7.

68. Mills, E.L., Pierce, K.A., Jedrychowski, M.P., Garrity, R., Winther, S., Vidoni, S., Yoneshiro, T., Spinelli, J.B., Lu, G.Z., Kazak, L., et al. (2018). Accumulation of succinate controls activation of adipose tissue thermogenesis. Nature 560, 102–106. 10.1038/s41586-018-0353-2.

69. Ryan, D.G., Murphy, M.P., Frezza, C., Prag, H.A., Chouchani, E.T., O’Neill, L.A., and Mills, E.L. (2019). Coupling Krebs cycle metabolites to signalling in immunity and cancer. Nat Metab 1, 16–33. 10.1038/s42255-018-0014-7.

70. Harber, K.J., de Goede, K.E., Verberk, S.G.S., Meinster, E., de Vries, H.E., van Weeghel, M., de Winther, M.P.J., and Van den Bossche, J. (2020). Succinate Is an Inflammation-Induced Immunoregulatory Metabolite in Macrophages. Metabolites 10, 372. 10.3390/metabo10090372.

71. Reddy, A., Bozi, L.H.M., Yaghi, O.K., Mills, E.L., Xiao, H., Nicholson, H.E., Paschini, M., Paulo, J.A., Garrity, R., Laznik-Bogoslavski, D., et al. (2020). pH-Gated Succinate Secretion Regulates Muscle Remodeling in Response to Exercise. Cell 183, 62–75.e17. 10.1016/j.cell.2020.08.039.

72. Mills, E.L., Harmon, C., Jedrychowski, M.P., Xiao, H., Garrity, R., Tran, N.V., Bradshaw, G.A., Fu, A., Szpyt, J., Reddy, A., et al. (2021). UCP1 governs liver extracellular succinate and inflammatory pathogenesis. Nat Metab 3, 604–617. 10.1038/s42255-021-00389-5.

73. Reddy, A., Winther, S., Tran, N., Xiao, H., Jakob, J., Garrity, R., Smith, A., Ordonez, M., Laznik-Bogoslavski, D., Rothstein, J.D., et al. (2024). Monocarboxylate transporters facilitate succinate uptake into brown adipocytes. Nat Metab 6, 567–577. 10.1038/s42255-024-00981-5.

74. Wilde, B.R., Chakraborty, N., Matulionis, N., Hernandez, S., Ueno, D., Gee, M.E., Esplin, E.D., Ouyang, K., Nykamp, K., Shuch, B., et al. (2023). FH variant pathogenicity promotes purine salvage pathway dependence in kidney cancer. Cancer Discov 13, 2072–2089. 10.1158/2159-8290.CD-22-0874.

75. Wu, Z., Bezwada, D., Cai, F., Harris, R.C., Ko, B., Sondhi, V., Pan, C., Vu, H.S., Nguyen, P.T., Faubert, B., et al. (2024). Electron transport chain inhibition increases cellular dependence on purine transport and salvage. Cell Metab 36, 1504–1520.e9. 10.1016/j.cmet.2024.05.014.

76. Tran, D.H., Kim, D., Kesavan, R., Brown, H., Dey, T., Soflaee, M.H., Vu, H.S., Tasdogan, A., Guo, J., Bezwada, D., et al. (2024). De novo and salvage purine synthesis pathways across tissues and tumors. Cell 187, 3602–3618.e20. 10.1016/j.cell.2024.05.011.

77. Ueno, D., Vasquez, J.C., Sule, A., Liang, J., van Doorn, J., Sundaram, R., Friedman, S., Caliliw, R., Ohtake, S., Bao, X., et al. (2022). Targeting Krebs-cycle-deficient renal cell carcinoma with Poly ADP-ribose polymerase inhibitors and low-dose alkylating chemotherapy. Oncotarget 13, 1054–1067. 10.18632/oncotarget.28273.

78. Nam, M., Xia, W., Mir, A.H., Jerrett, A., Spinelli, J.B., Huang, T.T., and Possemato, R. (2024). Glucose limitation protects cancer cells from apoptosis induced by pyrimidine restriction and replication inhibition. Nat Metab 6, 2338–2353. 10.1038/s42255-024-01166-w.

79. Zhao, X.H., Han, M.M., Yan, Q.Q., Yue, Y.M., Ye, K., Zhang, Y.Y., Teng, L., Xu, L., Shi, X.-J., La, T., et al. (2025). DNA replication stress underpins the vulnerability to oxidative phosphorylation inhibition in colorectal cancer. Cell Death Dis 16, 1–12. 10.1038/s41419-025-07334-4.

80. Trott, O., and Olson, A.J. (2010). AutoDock Vina: improving the speed and accuracy of docking with a new scoring function, efficient optimization, and multithreading. J Comput Chem 31, 455–461. 10.1002/jcc.21334.

81. O’Boyle, N.M., Banck, M., James, C.A., Morley, C., Vandermeersch, T., and Hutchison, G.R. (2011). Open Babel: An open chemical toolbox. J Cheminform 3, 33. 10.1186/1758-2946-3-33.

82. Ruiz-Ramos, A., Lallous, N., Grande-García, A., and Ramón-Maiques, S. (2013). Expression, purification, crystallization and preliminary X-ray diffraction analysis of the aspartate transcarbamoylase domain of human CAD. Acta Crystallogr F Struct Biol Cryst Commun 69, 1425–1430. 10.1107/S1744309113031114.

83. Studier, F.W. (2005). Protein production by auto-induction in high density shaking cultures. Protein Expr Purif 41, 207–234. 10.1016/j.pep.2005.01.016.

84. Prescott, L.M., and Jones, M.E. (1969). Modified methods for the determination of carbamyl aspartate. Analytical Biochemistry 32, 408–419. 10.1016/S0003-2697(69)80008-4.

85. Readshaw, J.J., Doyle, L.A., Puiu, M., Kelly, A., Nelson, A., Kaiser, A.J., McGuire, S.F., Peralta Acosta, J., Smith, D.L., Stoddard, B.L., et al. (2025). PglZ from Type I BREX phage defence systems is a metal-dependent nuclease that forms a sub-complex with BrxB. Nucleic Acids Res 53, gkaf540. 10.1093/nar/gkaf540.

86. Freie, B., Ibrahim, A.H., Carroll, P.A., Bronson, R.T., Augert, A., MacPherson, D., and Eisenman, R.N. MAX inactivation deregulates the MYC network and induces neuroendocrine neoplasia in multiple tissues. Sci Adv 11, eadt3177. 10.1126/sciadv.adt3177.

